# Circular RNA Circ-Cdr1as modulates Macrophage phenotype and Cardiac Reparative Function by Circ-Cdr1as-miR-7-Klf4 pathway

**DOI:** 10.1101/2025.02.21.639391

**Authors:** Carolina Gonzalez, Maria Cimini, Vandana Mallaredy, Cindy Benedict, Darukeshwara Joladarashi, Charan Thej, Zhongjian Cheng, May Trungcao, Amit Kumar Rai, Venkata Naga Srikanth Garikipati, Raj Kishore

## Abstract

**Background:** Mechanisms of macrophage switching from pro-inflammatory to anti-inflammatory phenotypes are not well understood. Circular RNAs (circRNAs), a new class of non-coding RNAs, are implicated in immune modulation. We recently identified circ-cdr1as as a regulator of macrophage phenotype in bone marrow derived macrophages (BMDM), however, their role in immunomodulation during cardiovascular injury remains unknown.

**Methods:** Cell-specific expression levels of circ-cdr1as was determined in a mouse hearts post-myocardial infarction (MI). Circ-cdr1as was overexpressed in fluorescently labeled BMDMs and injected into the ischemic myocardium immediately following MI. Effect of AAV9-mediated systemic delivery of circ-Cdr1as on post-MI cardiac function and structure was determined. Downstream mechanisms were studied using gain and loss of function strategies.

**Results:** Cardiac cell specific expression analysis showed significant downregulation of circ-cdr1as only in macrophages and cardiomyocytes. Overexpression of circ-cdr1as in BMDMs, injected into the ischemic myocardium retained their anti-inflammatory phenotype and significantly improved left ventricular (LV) functions and reduced infarct size. Systemic delivery of AAV9-circ-cdr1as showed similar cardiac reparative activity. Mechanistically, circ-cdr1as directly binds and sponge microRNA-7 and increases the expression of target KLF4. Loss and gain of function studies show that modulation of miR-7 and KLF recapitulates macrophage phenotypic changes.

**Conclusions:** Circ-cdr1as plays a crucial role in regulating the anti-inflammatory phenotype of macrophages through modulation of miR-7 and its target gene KLF4. Therefore, circ-cdr1as holds potential as an anti-inflammatory regulator in tissue inflammation post-cardiac injury.

**Novelty and Significance:** *What is Known?:*

- Despite continued research in elucidating mechanisms involved in cardiovascular disease and benefits of approved guideline-based therapies, the leading cause of deaths worldwide continues to be cardiovascular diseases (CVDs) with an increasing incidence of heart failure.
- Advances in high-throughput RNA sequencing (RNA-seq) allowed the identification of novel transcripts such as microRNAs (miRNA), long non-coding RNAs (lncRNAs), and circular RNAs (circRNA). circular RNAs have recently emerged as promising candidates for targeted therapy due to their circular structure that confers resistance to exonucleases, their capability to regulate gene expression by modulating miRNA activity, sequester proteins by acting as protein sponges,
- Several studies identified circRNAs to be differentially expressed following myocardial infarction (MI) and to play a role in immunity by contributing to the process of macrophage polarization, appropriate activation of macrophages when exposed to LPS, and inhibition of macrophage biogenesis. However, there are currently no published studies into the role of circular RNAs in the regulation of macrophage plasticity during cardiac injury.

*What New information Does This Article Contribute?:*

- We provide evidence that circ-cdr1as expression is downregulated in the heart 3 days post MI and specifically in cardiomyocytes and macrophages.
- Our study provides promising evidence that overexpression of circ-cdr1as may be cardioprotective by reducing cardiomyocyte apoptosis, enhancing angiogenesis, limiting infarct size, increasing percentage of anti-inflammatory macrophages, and overall preserving post-MI cardiac function.
- Mechanistically, we identified a reciprocal relationship between circ-cdr1as and miR-7 at 3 days post-MI and in naïve, pro-, and anti-inflammatory macrophages indicating circ-cdr1as role as a miRNA sponge. This suggests that circ-cdr1as/miR-7/Klf4 play a crucial role in cardiac injury and macrophage phenotype.

## Introduction

Cardiovascular disease (CVD) is the leading cause of morbidity and mortality, accounting for more than 900,000 deaths in the United States annually and despite advances in treatment and care, the total case numbers and long-term prognosis remain poor ^1^. Myocardial infarction (MI) is one of the most common forms of acute cardiac injury, resulting in the death of myocardial tissue and persistent inflammation ^2^.

The link between heart failure and inflammation was first described in 1990 by Levine et al., who reported elevated levels of tumor necrosis factor in heart failure patients with a reduced ejection fraction ^3^. Numerous studies thereafter corroborated this finding, revealing elevated levels of inflammatory mediators in patients experiencing acute decompensated heart failure and heart failure patients with preserved ejection fraction, suggesting a consistent inflammatory response present in all the manifestations of heart failure ^4–6^. In the initial stages of cardiac injury, dying cardiomyocytes release damage associated molecular patterns and pro-inflammatory cytokines to recruit leukocytes and initiate an inflammatory response. Neutrophils are the first leukocyte to be recruited, they release matrix degrading enzymes and reactive oxygen species. Macrophages are then recruited in high numbers to the infarcted myocardium and undergo phenotypic switching from pro-inflammatory phenotype in the early stage to anti-inflammatory phenotype in the later stage to control the initiation and resolution of the inflammatory response ^6–8^. Apart from their acknowledged function in the immune response, macrophages participate in crosstalk with other cardiac cells such as cardiomyocytes, fibroblast, immune cells, and endothelial cells to dictate the inflammatory and repair processes ^8^. As a result of macrophages being central regulators for the immune system, there have been attempts to target them via reduction, depletion, or impairment of mobilization and function. However, these strategies prolonged cardiac inflammation leading to insufficient clearance of dead cells and impaired tissue remodeling ^9–11^. Therefore, further investigation is needed to identify mechanisms involved in the modulation of the inflammatory response and their potential for therapeutic intervention. In recent years, non-coding RNAs (ncRNAs) have been investigated as regulatory RNAs involved in pathological process in CVD ^12^. In particular, circular RNAs have begun to garner interest as potential biomarkers and regulators of cardiac dysfunction ^13–16^ and immune response in infections, auto-immune diseases, and cancer ^17–20^.

Circular RNAs are newly discovered non-coding RNA generated from protein-coding genes with covalently closed loop generated by the pre-mRNA splicing machinery that backsplices to join a downstream 5’ splice site to an upstream 3’ splice site ^21,22^. The unique characteristic to lack free terminals allow circRNAs to be resistant to exonuclease degradation resulting in high stability. Studies have reported that circRNAs exhibit biological functions, including transcriptional and translational regulation, sequesters of miRNA and RBPs, and cell-to-cell communication ^14,23–26^. Circular RNAs are abundant in the eukaryotic transcriptome and have been identified to be differentially expressed during cardiovascular development and disease ^15,16^. Additionally, circRNAs have also been implicated in immune regulation ^27^. Notably, our previous findings identified circ-cdr1as to modulate phenotypic switching of macrophages to a more anti-inflammatory phenotype and overexpression of circ-cdr1as increased the transcription of anti-inflammatory markers in both pro- and anti-inflammatory macrophages *in vitro* ^28^. However, there are currently no reports on the role of circRNAs in macrophage biology and function during cardiac ischemia. Therefore, we aimed to understand the role of circ-cdr1as in macrophage function post-cardiac injury. In this current study, we provide evidence that either AAV9 mediated cardiac overexpression of circ-cdr1as and injection of BMDM overexpressing circ-cdr1as directly injected into the ischemic myocardium can increase anti-inflammatory macrophages in the injured myocardium, enhance angiogenesis and attenuate LV dysfunction post-MI. Mechanistically, we provide supporting evidence that circ-cdr1as may invoke physiological changes in macrophages by acting as a sponge for miRNA-7 to prevent the inhibition microRNA-7 target gene Klf4.

## Methods

A detailed description of all materials and methods can be found in the Online Supplemental Material. The data that support the findings of this study are available upon reasonable request.

### Experimental Animal Models

All animal procedures were performed following the approved protocols of the Institutional Animal Care and Use Committee (IACUC) of Temple University and conforms to the Guide for the Care and Use of the laboratory animals published by the US National Institutes of Health. Eight weeks old wild type (WT) male mice of C57BL/6 background and C57BL/6-Tg (UBC-GFP)30Scha/J with a C57BL/6 background were procured from Jackson Research Laboratory (Bar Harbor, ME) and randomly assigned to the experimental groups. Briefly, UBC-GFP transgenic mice express green fluorescent protein (GFP) directed by the human ubiquitin C promoter and allows *in vivo* leukocyte tracking based on distinct expression levels of GFP ^29,30^.

### Myocardial Infarction Surgery

MI was induced by permanent left anterior descending artery (LAD) ligation as we published before^40^. Briefly, a left intercostal thoracotomy was performed, and the ribs were retracted with 5-0 polypropylene sutures to open the chest. After the pericardium was opened, the left anterior descending artery (LAD) was ligated distal to the bifurcation between the LAD and diagonal branch using 8-0 polypropylene sutures through a dissecting microscope. After positive end-expiratory pressure was applied to inflate the lung fully, the chest was closed with 7-0 polypropylene sutures. A 22 G syringe was used to evacuate air from the chest cavity. The mice in the sham group underwent the same procedure except for the LAD ligation.

For the macrophage cell therapy study, immediately after LAD ligation, each mouse received intramyocardial injection of ∼500,000 AAV2 circ-cdr1as macrophages or AAV2 vehicle macrophages or saline. For the second *in vivo* study, mice received 1×10^11^vp/mL of AAV9 circ-cdr1as by tail vein injection 14days prior to MI as previously described ^31–34^.

### Circ-cdr1as Expression Plasmid and AAV Construction

Our circ-cdr1as expression plasmid was generated as previously described ^28^. Briefly, we used pcDNA3.1 (+) Lacasse2 MCS exon vector (Addgene plasmid # 69893; http://n2t.net/addgene:69893; RRID:Addgene_69893) ^35^ with fragment insertion in *PacI* and *SacII* sites of pcDNA3.1 (+) Lacasse2 MCS exon vector (Figure S2A). For *in vivo* expression, the sequence was cloned into a self-complementary AAV backbone plasmid (pTRUFR) and expression of the circRNA is driven by a CMV promoter and flanked by AAV9 ITRs (*in vivo* tail vein injections) or AAV2 (*ex vivo* overexpression in macrophages). AAV9-GFP and AAV2-empty vector were used as controls. All viruses were produced using triple transfection technique. ^36–38^

### Left Ventricle Heart Tissue Collection and Cardiac Cell Isolations

To isolate RNA from the LV tissue, the heart was quickly removed from the chest and perfused with phosphate buffered saline (PBS) 3x. The LV tissue was excised and minced prior to RNA extraction. Adult mouse cardiomyocytes, endothelial cells, and fibroblast were isolated as previously described ^39^.

### Echocardiography

Transthoracic two-dimensional M-mode echocardiography using the Vevo 2100 equipped with 30 MHz transducers (VisualSonics, Toronto, ON, Canada) was performed before MI (baseline), and at 7, 21, and 28 days after surgery as described previously ^40^. Mice were anesthetized with a mixture of 1.5% isoflurane and oxygen (1 L/min) with an isoflurane delivery system (Viking Medical, Medford, NJ).

The internal diameter of the LV was measured in the short-axis view from M-mode recordings; percent ejection fraction (% EF) and fractional shortening (% FS) were calculated using corresponding formulas as previously described ^40^.

### FACS Analysis of Cardiac and Bone Marrow Derived Macrophages

FACS analysis of cardiac and bone marrow derived macrophages was performed as described in our earlier publication^28^.

### Cell Culture and in vitro Studies

Bone marrow (BM) monocytes were isolated from bone marrow mononuclear cells of C57BL/6 male (approximately 8–10-week-old) mice by density-gradient centrifugation with histopaque-1083 (Sigma) and red blood cells were removed with NH_4_Cl (Stemcell Technologies) as previously described ^28,41^. The monocyte population was cultured in RPMI 1640 1× (Gibco) with 20% FBS, 1% penicillin-streptomycin solution, and 20% of L929-conditioned medium. The media was changed the day after culture, on day 3, and on day 5–7. By day 5–7, high purity of macrophages can be observed as previously noted ^42^. On day 5–7, change to fresh stimulation media: for pro-inflammatory activation ^43^, 100 ng/mL IFNγ (R&D) and 100 ng/mL TNFα (PromoKine); for anti-inflammatory activation ^44^, 10 ng/mL IL-4 (R&D), 10 ng/mL IL-10 (R&D), and 20 ng/mL TGF-β (R&D) for a period of 24 h as previously described ^28^. The Raw 264.7 monocytes were cultured in DMEM-F12 medium (Corning, USA) with 10% fetal bovine serum (FBS) and 1% penicillin-streptomycin solution. Raw 264.7 cells were activated with 100 ng/mL of LPS for 24 hours. BMDMs or RAW 264.7 cells were transfected for 24 hours with 100nM miRNA mimic miR-7b-3p (Assay ID MC26348, Thermofisher) or anti-miR-7b-3p (Assay ID: MH26348, Thermofisher) or corresponding controls (mirVana™ miRNA Inhibitor/Mimic, Negative Control #1, Thermofisher). Lipofectamine RNAiMAX (Thermofisher) was used to facilitate transfection. The cells were transfected prior to polarization for 24 hours. For knockdown of circ-cdr1as, BMDMs were transfected with MOI 15 lentiviral shRNA (titer concentration for shRNA scrambled: 2.9 × 10^9^; shRNA circ-cdr1as: 3.67 × 10^9^ vp/mL) as previously described ^28^. For overexpression of Klf4, BMDMs were transfected with Klf4 (NM_010637) Mouse Tagged ORF Clone Lentiviral Particle (Origene) (titer concentration 1.7×10^7 TU/mL). Lentivirus Lenti ORF control particles of pLenti-C-mGFP-P2A-Puro were used as a negative control (Origene) (titer concentration 1.0×10^7 TU/mL). For knockdown of Klf4, BMDMs were transfected with Klf4 Mouse shRNA Lentiviral Particle generated from Origene using four unique 29-mer target-specific shRNA (titer concentration 1.7×10^7 TU/mL) and one scramble control (titer concentration 1.0×10^7 TU/mL)., all expressing GFP. Briefly, the cells were transfected with lentivirus and viral enhancer (ABM) for 48 h. The media was then changed to media containing 3μg/mL of puromycin antibiotic for selection of stably transfected cells. The cells were polarized for 24 h. Cells were harvested and changes in the expression or transcription of macrophage markers were measured by FACS or RT-qPCR, respectively.

### Ex vivo Overexpression of circ-cdr1as in BMDMs

BMDMs were transfected with 500 MOI of AAV2 circ-cdr1as (titer 4×10^9^ vp/mL). The cells were transfected with AAV2 and viral enhancer (VirallentryTM ABM). The following day, the media was changed to media containing 3 μg/mL of G418 antibiotic for selection of stably transfected cells. Cells were harvested and circ-cdr1as overexpression was verified by RT-qPCR.

### circRNA Pull Down Assay

Circ-cdr1as was captured using magnetic instant capture beads with circ-cdr1as single probe covalently attached to the surface of nanomagnetic particles designed by ElementZero Biolabs to target only the backsplicing junction. This allows the RNA affinity purification with magnetic instant capture beads for the capture of RNA-miRNA or RNA-protein complexes from chemically cross-linked material. The following protocol was used according to manufacturer instructions. MagIC-beads-RNA-enrichment-protocol-proteome-capture (elementzero.bio).

### Statistical Analysis

Data are expressed as mean ± standard error of the mean (S.E.M.) represent at least 3 independent biological experiments. Data visualization and statistical analyses were performed using GraphPad Prism 10.4.1 software (GraphPad, La Jolla, CA). Mean of the groups were compared using unpaired two tail t-test (for two groups) and One-way ANOVA (more than two groups) with Tukey’s multiple comparisons test. P values of <0.05 indicate statistical significance.

## Results

### Circular RNA cdr1as is Downregulated in Post-MI Hearts

We previously profiled circRNAs expression in bone marrow-derived macrophages polarized to either pro-inflammatory or anti-inflammatory phenotype and identified circ-cdr1as as a regulator of anti-inflammatory, M2-like macrophage phenotype ^28^. Therefore, we investigated circ-cdr1as expression at different time points post-MI. Circ-cdr1as was significantly downregulated at 3-5 days post-MI, the time during which pro-inflammatory macrophages are most prominent and upregulated starting at 7 to 28 days post-MI, the time in which inflammation subsides (Figure 1A). We then assessed circ-cdr1as expression in specific cardiac cell types in the post-MI left ventricular tissue. Circ-cdr1as expression was significantly downregulated in macrophages and cardiomyocytes. However, circ-cdr1as expression was not affected in endothelial cells or fibroblast (Figure 1B). These results suggest cell-specific downregulation of circ-cdr1as after MI, particularly in macrophages and cardiomyocytes.

**Figure 1.**
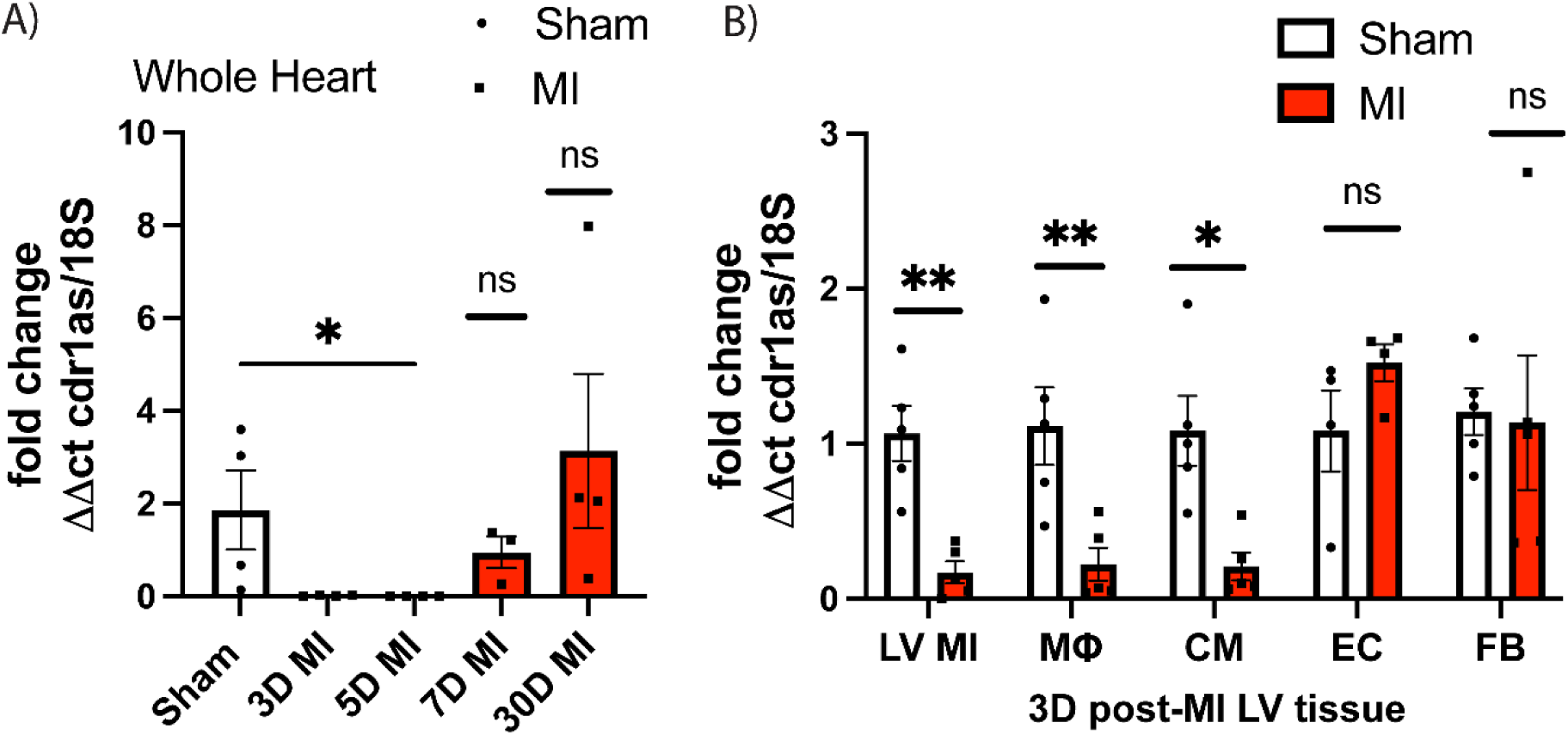
Circular RNA cdr1as expression in sham and MI hearts. (A) qRT-qPCR of circ-cdr1as expression at different time points in post-MI hearts compared to sham, normalized to 18S. N=3-4/group. (One-way ANOVA). (B) qRT-PCR analysis of circ-cdr1as expression in 3D post-MI left ventricular (LV) tissue including isolated macrophages (MΦ) (Aria sorted for F4/80+ cells), cardiomyocytes (CM), endothelial cells (EC) (CD31+ cells), and fibroblast (FB) cells isolated from 3D post-MI tissue compared to sham controls, normalized to 18S. (Two-sided unpaired t-test). N-4=5/group. Data are mean ± SEM. NS, non-significant, *p<0.05, ** p<0.01. MΦ, macrophages; CM, cardiomyocytes; EC, endothelial cells; FB, fibroblast.

### Circ-cdr1as overexpressing Bone Marrow Derived Macrophages Retain their M2-like Macrophage Phenotype when Injected in Post-MI Hearts

We utilized C57BL/6-Tg (UBC-GFP)30Scha/J with a C57BL/6 background to isolate GFP+ BMDMs to track macrophages in the myocardium after injection. GFP-BMDM were ex vivo transduced with AAV2 control and AAV2 circ-cdr1as vector and 5×10^5^ cells were injected into the MI border zone immediately LAD ligation. We performed FACS analysis of the percentage of GFP+ cells in mice 5 days post-MI (Figure 2A). The percent of GFP+ cells between AAV2 vehicle MΦ and AAV2 circ-cdr1as MΦ were comparable (Figure 2A). We further investigated the percentage of GFP/CD206+ cells and GFP+CD206+ cells normalized to gram of tissue in the border zone/infarcted region of these groups and found that AAV2 circ-cdr1as MΦ had a significant percentage of cells that were GFP/CD206+ compared to AAV2 vehicle MΦ group (Figure 2C). We then examined the ratio of Ly6c+ (pro-inflammatory macrophage marker) or CD206+ (anti-inflammatory macrophage marker) cells. AAV2 circ-cdr1as MΦ group had the highest percentage of CD206+ cells compared to Ly6c+ cells (Figure 2B, Figure S1B). We validated our FACS analysis with immunohistochemistry staining for GFP and CD206 at 5 days post-MI. AAV2 circ-cdr1as MΦ group had the highest percentage of GFP/CD206+ cells (Figure 2D, E). Additionally, we also stained with CD45 (leukocyte marker) and CD163 (anti-inflammatory macrophage marker) and found an increase in CD163+ cells in AAV2 circ-cdr1as MΦ group compared to saline and AAV2 vehicle MΦ groups (Figure S1C, D). Taken together, these results demonstrate that injection of AAV2 circ-cdr1as MΦs directly into the injured heart promotes more anti-inflammatory macrophages and that these cells can maintain an M2-like phenotype in the initial stages of cardiac injury, during which inflammation is high.

**Figure 2.**
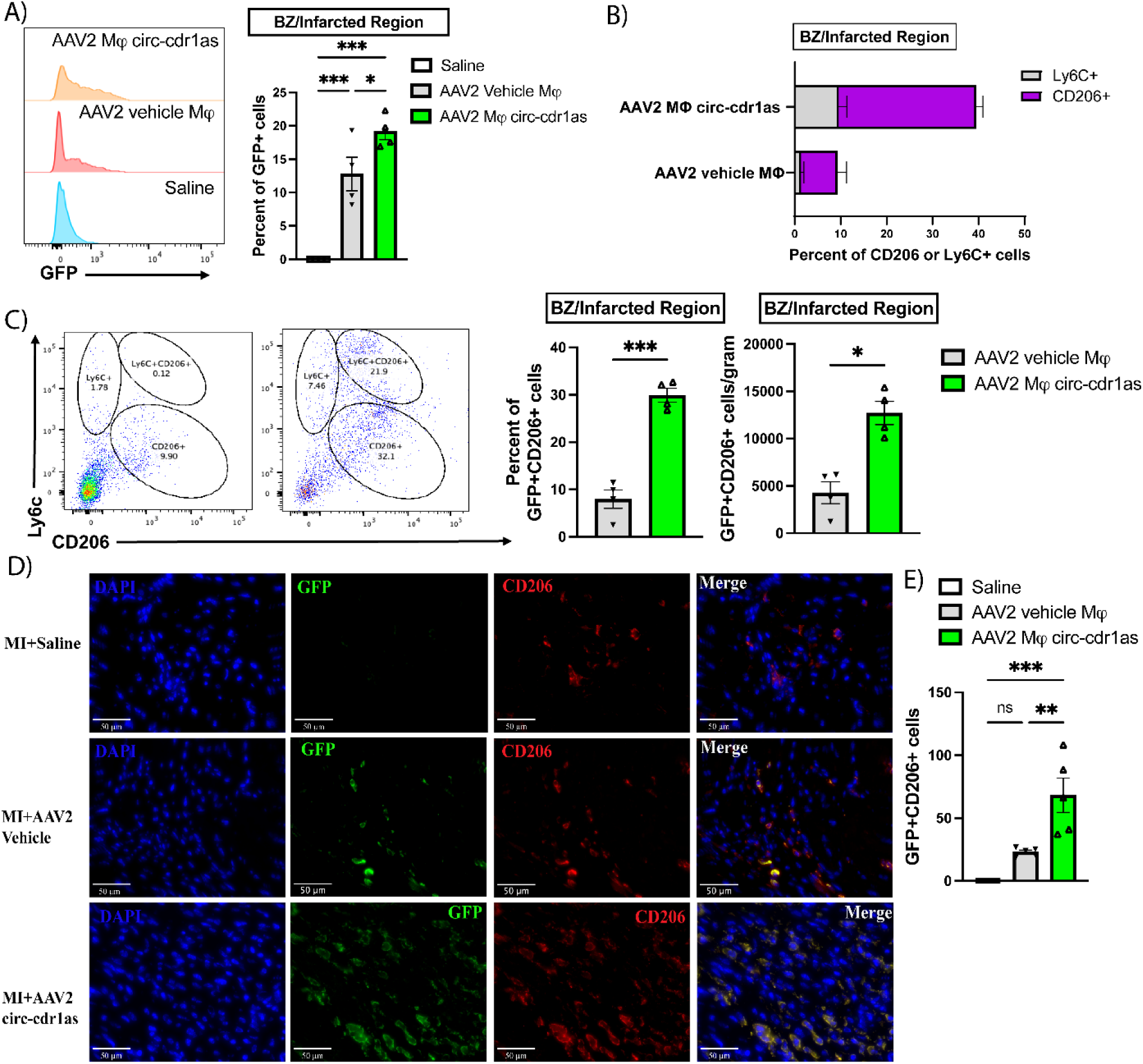
GFP+ circ-cdr1as overexpressing macrophages retain M2-like phenotype in ischemic heart. (A) FACS analysis of %GFP+ cells in the LV MI region at 5D post-MI of all groups. N=4/group. (B) FACS analysis of %GFP/CD206+ cells in the LV MI region at 5D post-MI of all. N=4/group. (C) FACS analysis of the percentage of CD206 or Ly6C cells found within the LV MI region 5D post-MI (left) and normalized cells/gram (right) for all groups. N=4/group (Two-sided unpaired t-test). (D) Representative photomicrographs (50µm) of GFP (green) and CD206 (red) positive cells and nuclei DAPI (blue)/40X. (E) Quantification of GFP/CD206+ cells. N= 5/group. Data are mean ± SEM. (One-way ANOVA). NS, non-significant, *p<0.05, ** p<0.01, *** p<0.001. GFP, green-fluorescent protein; Ly6C, M1 macrophage marker; CD206, M2 macrophage marker.

### Injection of circ-cdr1as Overexpressing BMDMs into the Ischemic Myocardium Improved LV Function and Reduced Infarct Size

Reduced levels of circ-cdr1as in the left ventricle 3 days post-MI, in macrophages, and in cardiomyocytes suggested a possible physiological role for circ-cdr1as in MI pathophysiology. Therefore, we tested if exogenous delivery of circ-cdr1as overexpressing macrophages at an early stage of cardiac injury could promote an anti-inflammatory response and thereby attenuate LV remodeling and dysfunction post-MI. We determined the effect of cell therapy on LV function by echocardiography at 7,21-, and 28-days post-MI (Figure 3B). Percent ejection fraction (EF) and fractional shortening (FS) at baseline and 7 days were similar in all groups (Figure 3B). Circ-cdr1as overexpressing macrophages significantly improved %EF and %FS at 21- and 28-days post-MI compared to saline and AAV2 MΦ vehicle (Figure 3B). Analysis of the LV end diastolic or systolic diameter indicated significant improvement with circ-cdr1as overexpressing macrophages treatment (Figure 3B). Additionally, heart weight was reduced in AAV2 circ-cdr1as MΦs group compared to AAV2 MΦ vehicle (Figure 3A). We next assessed infarct size by Masson’s trichrome staining at 28 days post-MI in mice and found a significant decrease in infarct area compared to AAV2 vehicle and saline groups (Figure 3C, D). In order to determine the effect of circ-cdr1as overexpressing macrophages on myocardial neovascularization, we stained sections with CD31 and α-SMA. We found increased capillary density and α-SMA+ arterioles in the border zone/infarcted area at 28 weeks post-MI in mice that received circ-cdr1as overexpressing macrophages compared to mice that received AAV2 vehicle macrophages or saline (Figure 3E, F).

**Figure 3.**
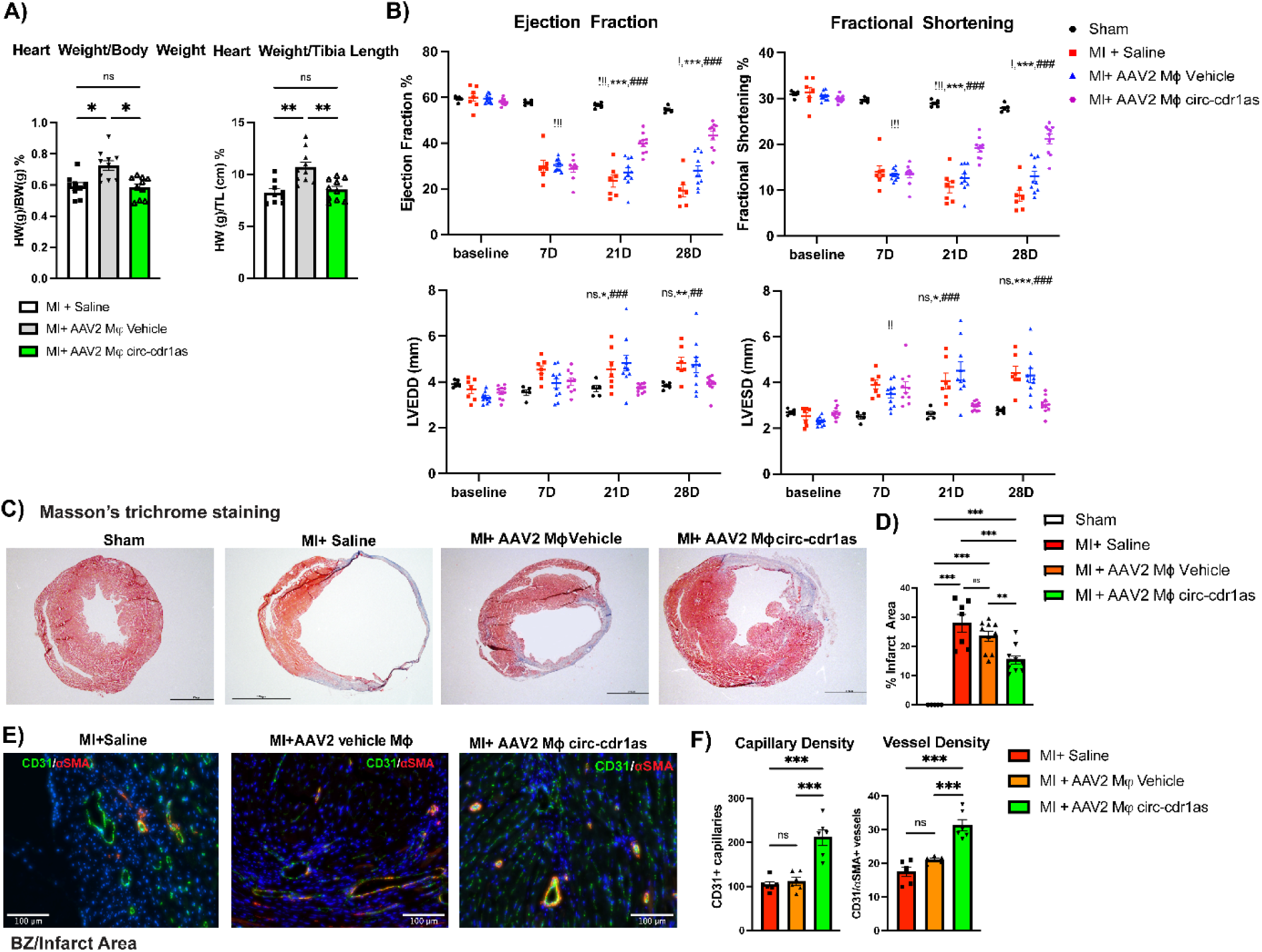
Circ-cdr1as overexpressing macrophages injected into the myocardium improved cardiac function and reduced infarct size 28 days post-MI in mice. (A) Heart weight/body weight (left) and Heart weight/tibia length (right) ratios in MI saline, AAV2 MΦ vehicle, and AAV2 circ-cdr1as MΦ groups. (B) GFP+ circ-cdr1as overexpressing macrophages improved LV function (% ejection fraction, % fractional shortening, LVEDD (mm), and LVESD (mm)) measured by echocardiography. (!: sham vs AAV2 circ-cdr1as MΦ; *: MI+ Saline vs AAV2 circ-cdr1as MΦ; # AAV2 MΦ vehicle vs AAV2 circ-cdr1as MΦ) N= 5 (sham), N= 7-10 (all other groups) (C) Representative Masson’s Trichrome stained heart sections in sham, saline, AAV2 MΦ vehicle, and AAV2 circ-cdr1as MΦ at 28 days post-MI. (D) Quantification of infarct size at 28D days post-MI in each group. N=5 (sham), N=7-10 (all other groups). ED) Representative images of CD31+ and αSMA+ cells in the border zone/infarcted area of the left ventricle at 28D post-MI. Capillaries were stained with CD31+ (green), arterioles stained with αSMA (red), and nuclei were stained with DAPI (blue). (F) Capillary density and vessel density were quantified as the number of CD31+ capillaries/20X or CD31/αSMA+ vessels/20X, respectively. N=6/group. One-way ANOVA. Data are mean ± SEM. NS, non-significant, *p<0.05, ** p<0.01, *** p<0.001.

### Administration systemic of AAV9 circ-cdr1as Enhanced Cardiac circ-cdr1as Expression and improved cardiac function after MI injury

In addition to macrophages, cell specific analysis (figure 1) also indicated post-MI downregulation of circ-cdr1as also in cardiomyocytes. We examined if cardiac overexpression of circ-cdr1as would improve cardiac function and remodeling. We utilized AAV9 vector, which has a higher tropism for cardiomyocytes, to deliver circ-cdr1as by intravenous injection. We injected 1×10^11^vp/mL by tail vein and examined the expression of circ-cdr1as in the LV tissue 14 days after tail vein injection and found a significant increase in the cardiac expression of circ-cdr1as (Figure S2C). We then verified increased circ-cdr1as expression at 5- and 28-days post-MI by RT-qPCR (Figure S2C). Although AAV9 has a higher tropism for cardiomyocytes and the highest expression of circ-cdr1as was found in the cardiomyocytes, since circ-cdr1as expression is driven by a CMV promoter, we also observed increased expression of circ-cdr1as in macrophages, endothelial cells, and fibroblast at 5 days post-MI, although to a lesser extent than cardiomyocytes (Figure S2D). We further validated cardiomyocyte tropism of AAV9 by intravenous injections of control AA9-GFP vector and verifying the expression of GFP in cardiomyocytes by immunohistochemistry (Figure S2E). Administration of AAV9 circ-cdr1as significantly improved cardiac LV function examined by echocardiography. The percent EF and FS were similar at baseline for AAV9-GFP and AAV9 circ-cdr1as groups. There was a significant improvement in %EF and %FS for the AAV9 circ-cdr1as group at 21- and 28-days post-MI compared to the AAV9 vehicle group. Additionally, the LV end diastolic and systolic diameters indicated significant restoration of LV dimension at 28 days with AAV9 circ-cdr1as treatment (Figure 4A). We also found a significant increase in myocardial neovascularization reflected by increased capillary density and α-SMA+ arterioles in the border zone and infarcted areas at 28 days post-MI in mice treated with AAV9 circ-cdr1as compared to AAV9-GFP (Figure 4B, C). We next examined the percentage of infarcted area between these groups by Masson’s trichrome staining at 28 days post-MI and found a significant reduction in the infracted area compared to the AAV9-GFPgroup (Figure 4D, E). To determine if cardiomyocyte size (cardiomyocyte hypertrophy) was affected by AAV9 circ-cdr1as treatment at 28 days post-MI, we stained with WGA and quantified the area/cell. We found no significant change in cardiomyocyte size in mice treated with AAV9 circ-cdr1as or AAV9-GFP (Figure 4F, G).

**Figure 4.**
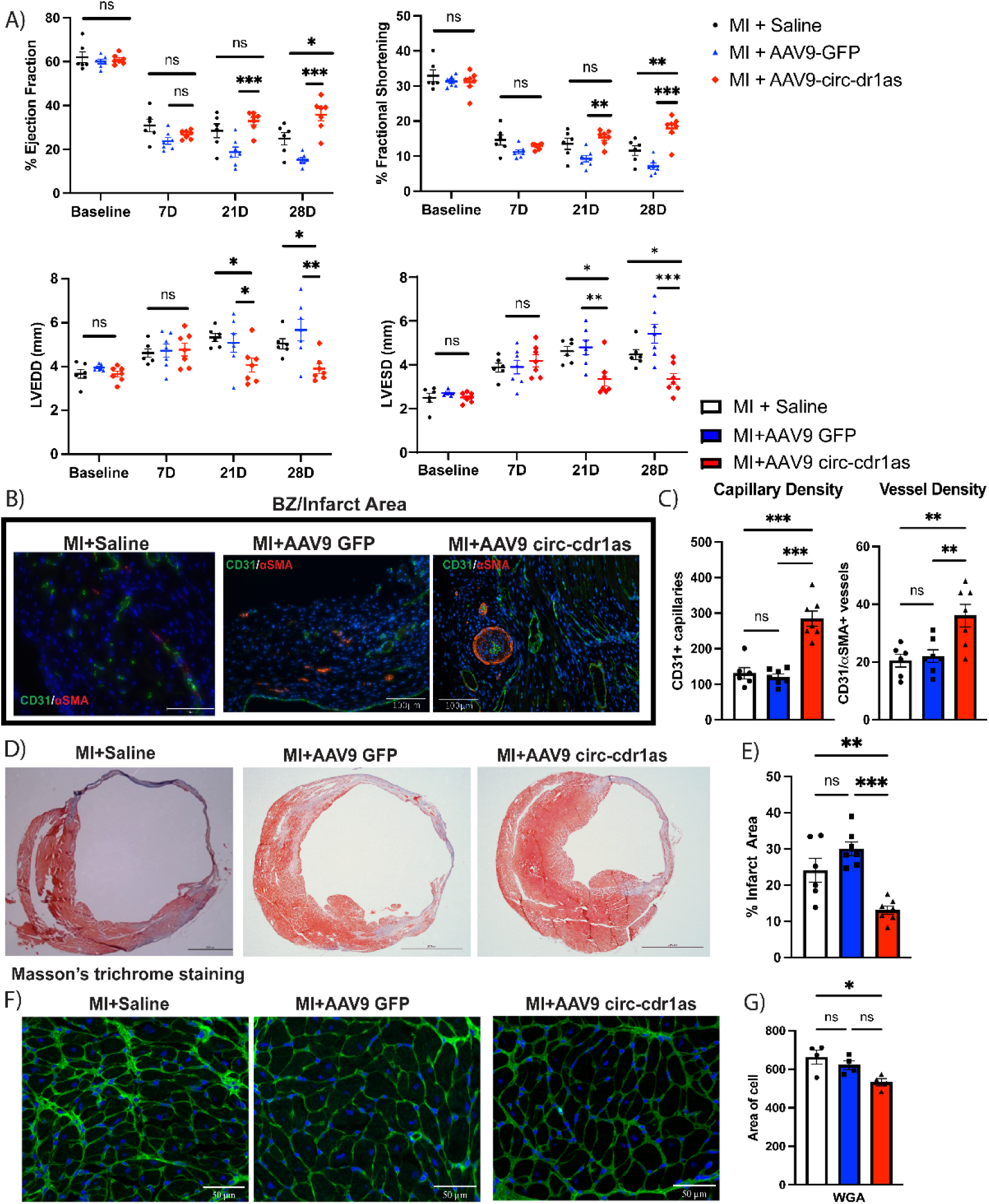
AAV9 circ-cdr1as overexpression improves cardiac function and reduces infarcted area in mice. (A) AAV9 circ-cdr1as significantly improved heart function at 28 days post MI (% ejection fraction, % fractional shortening, LVEDD (mm), and LVESD (mm)) measured by echocardiography compared to saline or AAV9-GFP. N=6-7/group. (B) Representative images (100µm) of capillary and vessel density measured in infarcted area/border zone area at 28 days post MI. Capillaries were stained with CD31 (green), arterioles stained with αSMA (red), and nuclei with DAPI (blue). (C) Capillaries were quantified as the number of CD31+ cells/20X and vessels were quantified as the number of CD31/αSMA + cells. N=6-7/group. (D) Representative images of Masson’s Trichrome stained heart sections in AAV9 GFP and AAV9 circ-cdr1as groups at 28 days post-MI and (E) quantitative analysis of infarct size. N=6-7/group. (F) Representative images of WGA (50µm) staining indicating cardiomyocyte hypertrophy and (F) quantification of the area/cell 40X. N=4/group. One-way ANOVA. Data are mean ± SEM. NS, non-significant,

### Cardiac Overexpression of circ-cdr1as Reduces Cardiomyocyte Apoptosis and Increases Anti-Inflammatory Macrophages 5 days post-MI

Anti-inflammatory macrophages exhibit a pro-regenerative phenotype and secrete high levels of anti-inflammatory cytokines including IL-10 and growth factors such as vascular endothelial growth factor (VEGF) and myeloid-derived growth factor known to be cardioprotective ^43,45^. Therefore, we assessed the role of circ-cdr1as on apoptosis of cardiomyocytes 5 days post-MI by TUNEL assay. We found a decrease in the percentage of cardiomyocyte apoptosis in mice treated with AAV9 circ-cdr1as compared to AAV9-GFP vehicle (Figure 5A). Interestingly, we found that treatment with AAV9 circ-cdr1as increased the number of anti-inflammatory macrophages present at the site of injury, indicated by increase in CD163 and CD206 positive cells 5 days post-MI (Figure 5B, C). We further validated these results by FACS analysis investigating the percentage of pro-(Ly6c) or anti-inflammatory (CD206) macrophages (Figure S3A). We found that AAV9 circ-cdr1as treatment increased the ratio of CD206+ cells in the LV tissue compared to AAV9 vehicle at 5 days post-MI (Figure 3A). Meanwhile, the percentage of total immune cells (CD45+ cells) remained comparable between groups (Figure S3B).

**Figure 5.**
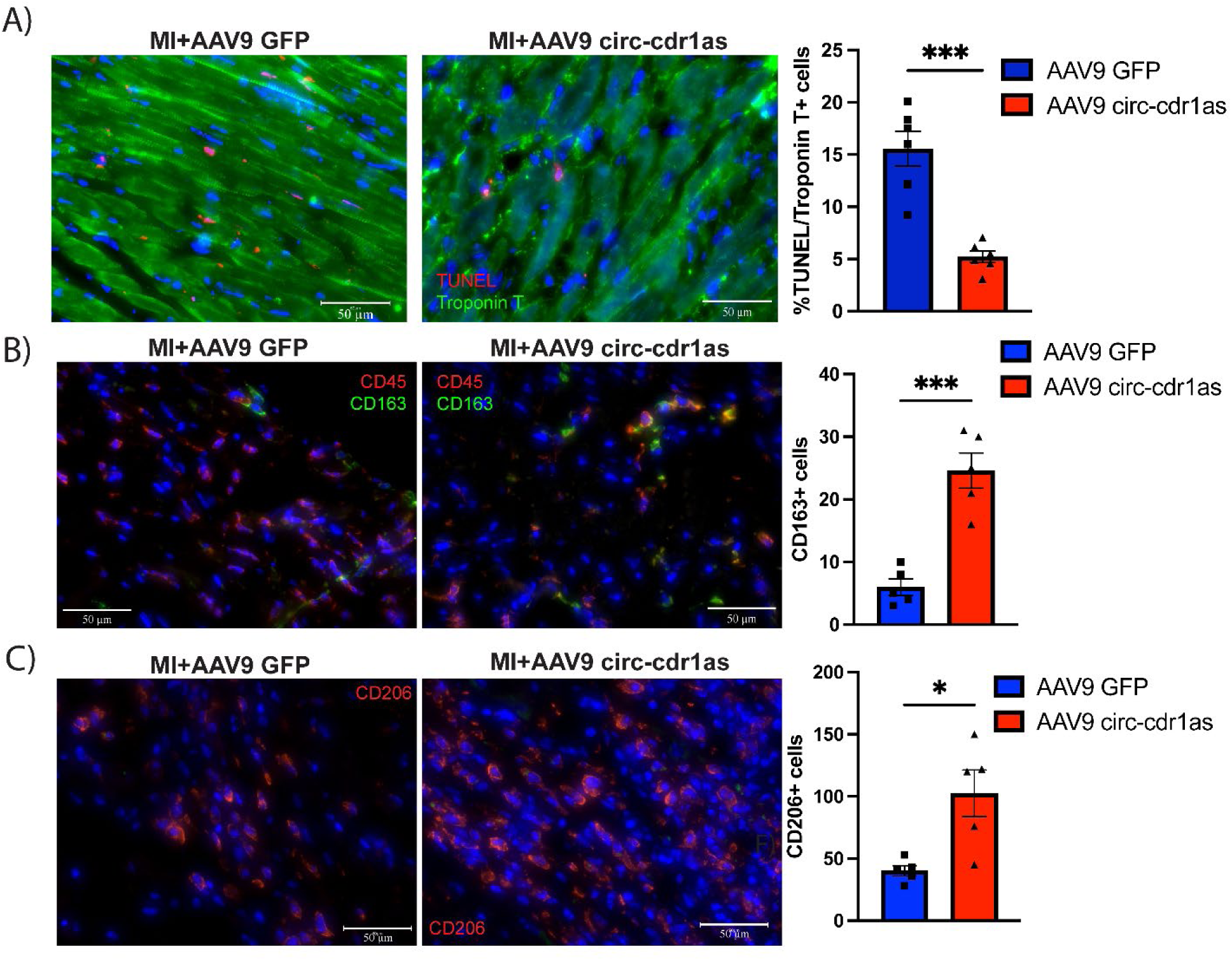
Cardiac overexpression of circ-cdr1as reduces cardiomyocyte cell death enhances anti-inflammatory macrophages 5 days post-MI. (A) Representative photomicrographs (50µm) of TUNEL (red)/Troponin T (green)staining in AAV9-GFP and AAV9 -circ-cdr1as treated mice and quantification of TUNEL/Troponin T + cells represented as the percent of TUNEL/Troponin T + cells and DAPI-stained nuclei/40X. N=6/group. (B) Representative photomicrographs (50µm) of CD163 (green) and CD45 (red) positive cells and quantification of CD163+ cells/40X. N=5/group (C) Representative photomicrographs (50µm) of CD206 (red) positive cells and quantification of CD206+ cells/40X. N=5/group. Unpaired t-test. Data are mean ± SEM. NS, non-significant, NS, non-significant, *p<0.05, ** p<0.01, *** p<0.001. GFP, green-fluorescent protein; Ly6C, M1 macrophage marker; CD206, M2 macrophage marker; CD45, leukocyte marker; CD163, M2 macrophage marker.

### Circular RNA cdr1as Modulates miR-7 Expression in Injured Hearts and BMDMs

Published literature including *in silico* analysis in our most recent paper identified miRNA-7 as a circ-cdr1as target based on Arraystar’s miRNA target prediction software utilizing TargetScan and miRanda (Figure S4B) ^28,46–51^. Circ-cdr1as is suggested to act as miR-7 sponge. Therefore, we investigated the expression of miR-7b-3p in the LV tissue, cardiomyocytes, and macrophages at 3 days post-MI. We identified a significant upregulation of miR-7b-3p in the LV tissue, cardiomyocytes and cardiac macrophages, at a time point when circ-cdr1as is significantly downregulated (Figure 6A). Based on these results, we performed a circ-cdr1as pull-down assay with probes specifically designed to target the circ-cdr1as backsplicing junction covalently attached to the surface of magnetic particles. Using RAW264.7 macrophage cells polarized to pro- or anti-inflammatory phenotype, circ-cdr1as pulldown assays confirmed its direct binding to miRNA-7b-3p (Figure 6B, Figure S4A). To determine if miR-7b-3p directly plays a role in BMDM polarization, we treated BMDMs with antagonist of miR-7 or miR-7 mimic or their respective controls (Figure S4C). FACS analysis demonstrated that treatment of miR-7 mimics increased percentage of pro-inflammatory marker CD86 (Figure 6C) and reduction in anti-inflammatory marker CD206 in naïve, pro-, or anti-inflammatory macrophages (Figure 6D, Figure S4E). Conversely, treatment with anti-miR-7 had the oppositive effect (Figure 6C, D, Figure S4E). Transcriptional levels of anti-inflammatory lineage markers (Arg-1 and CD206) were upregulated, and transcriptional levels of pro-inflammatory markers (Ly6C and INOS) were downregulated in BMDMs treated with anti-miR-7. Meanwhile, treatment with miR-7 indicated the opposite effect (Figure S4D). These findings suggest that circ-cdr1as plays a role in regulating miR-7 that results in changes to polarization of macrophage phenotype.

**Figure 6.**
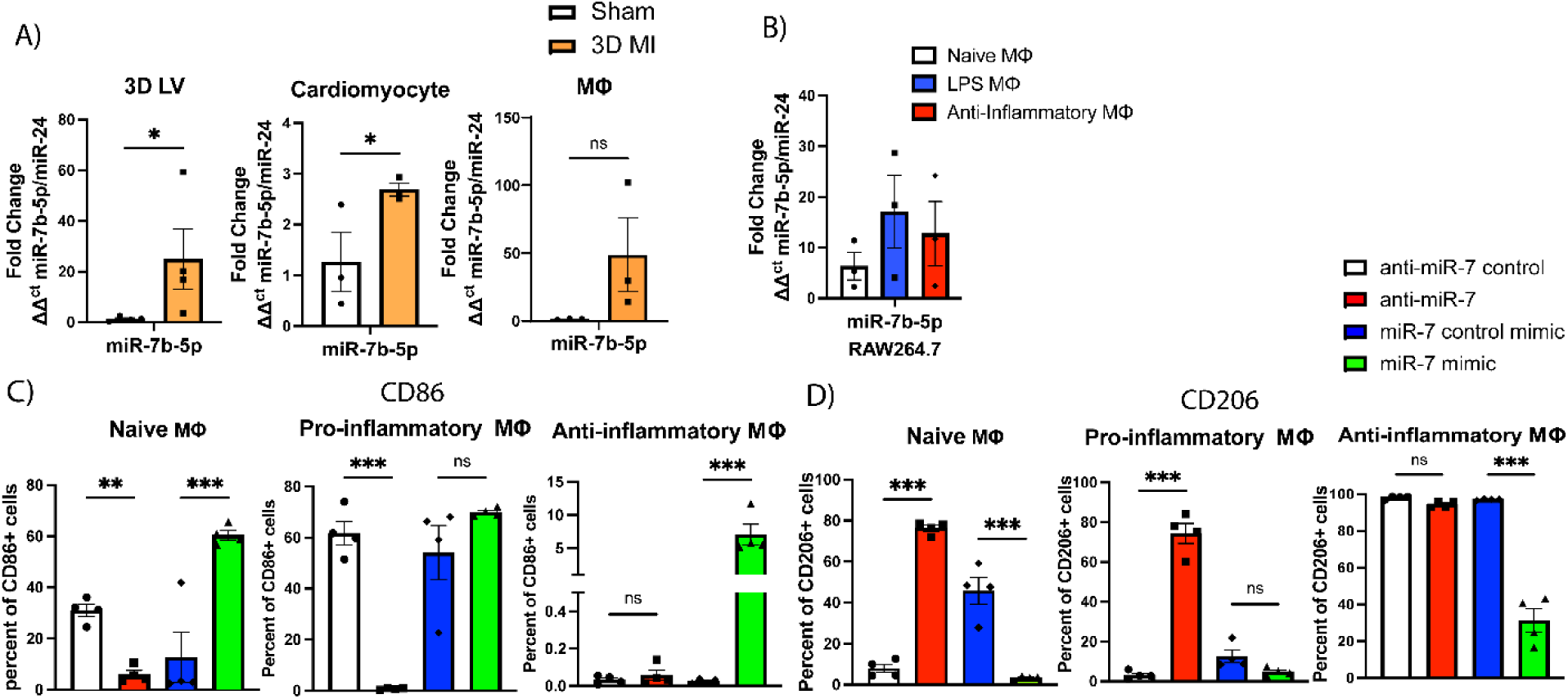
Inverse relationship between miR-7b-5p and circ-cdr1as in injured hearts and macrophages. (A) qRT-qPCR of miR-7b-3p expression in 3D post-MI left ventricular (LV) tissue including isolated macrophages (MΦ) (Aria sorted for F4/80+ cells), cardiomyocytes (CM) compared to sham controls, normalized to miR-24. (Two-sided unpaired t-test). N=3-4/group. (B) Level of miR-7b-5p after pull-down assessed by RT-qPCR in RAW 264.7 cells unpolarized or polarized with LPS or anti-inflammatory cytokines. (One-Way ANOVA). N= 3/group. (C) FACS analysis of F4/80/CD86 + cells (pro-inflammatory MΦ marker) or (D) F4/80/CD206+ cells (anti-inflammatory MΦ marker) in naïve, pro-inflammatory, and anti-inflammatory macrophages treated with miR-7 mimic or anti-miR and their respective controls. N= 4/group (One-way ANOVA). Data are mean ± SEM. NS, non-significant, *p<0.05, ** p<0.01, *** p<0.001. MΦ, macrophages; CM, cardiomyocytes; pro-inflammatory markers: Ly6C, lymphocyte antigen 6 family member C1; INOS, inducible nitric oxide synthase; anti-inflammatory markers: CD206; Arg-1, arginase 1.

### MicroRNA-7 target Klf4 is Upregulated in BMDMs Treated with Anti-miR-7

Recent evidence has shown that a target of miRNA-7 is Krüppel-like factor 4 (Klf4). Klf4 is associated with inflammatory responses and was indicated to be upregulated when miR-7 was inhibited ^52^. Thus, to explore the possible connection between miR-7 and Klf4 in macrophages, we assessed the comparative expression levels of Klf4 in the heart three days after myocardial infarction, in BMDMs subjected to circ-cdr1as overexpression or knockdown (by shRNA circ-cdr1as), and in BMDMs treated with anti-miR-7 or miR-7 mimic. We observed downregulation of Klf4 concomitant to increased miR-7 expression in LV tissue 3 days post-MI (Figure 7A). Additionally, Klf4 was significantly upregulated during cardiac overexpression of circ-cdr1as compared to the control (Figure 7B). Naïve, pro-, and anti-inflammatory macrophages had an upregulation of Klf4 expression when BDMDs overexpressed circ-cdr1as and a downregulation when circ-cdr1as was knocked down (Figure 7C). We saw the opposite relationship with miR-7 and Klf4, its expression was upregulated in Naïve, pro-, and anti-inflammatory macrophages when miR-7 was inhibited and a downregulated when BMDMs were treated with miR-7 mimic (Figure 7D). Gain and loss of klf4 experiments showed that overexpression of Klf4 in BMDMs upregulated percent of CD206+ cells, while knockdown of Klf4 upregulated percent of CD86+ cells (Figure 7E, Figure S5B). Transcriptional levels of anti-inflammatory lineage markers (Arg-1 and CD206) were upregulated, and transcriptional levels of pro-inflammatory markers (Ly6C and INOS) were downregulated BMDMs overexpressing Klf4. On the contrary, the knockdown of Klf4 indicated the opposite effect (Figure S7C, D). We also utilized kenpaullone, a small molecule that functions as a CDK1/GSK-3β inhibitor known to inhibit Klf4 with similar results ^53–55^. Collectively, our data indicates that circ-cdr1as modulates miR-7 to prevent interaction with and expression of Klf4 to alter macrophage phenotype.

**Figure 7.**
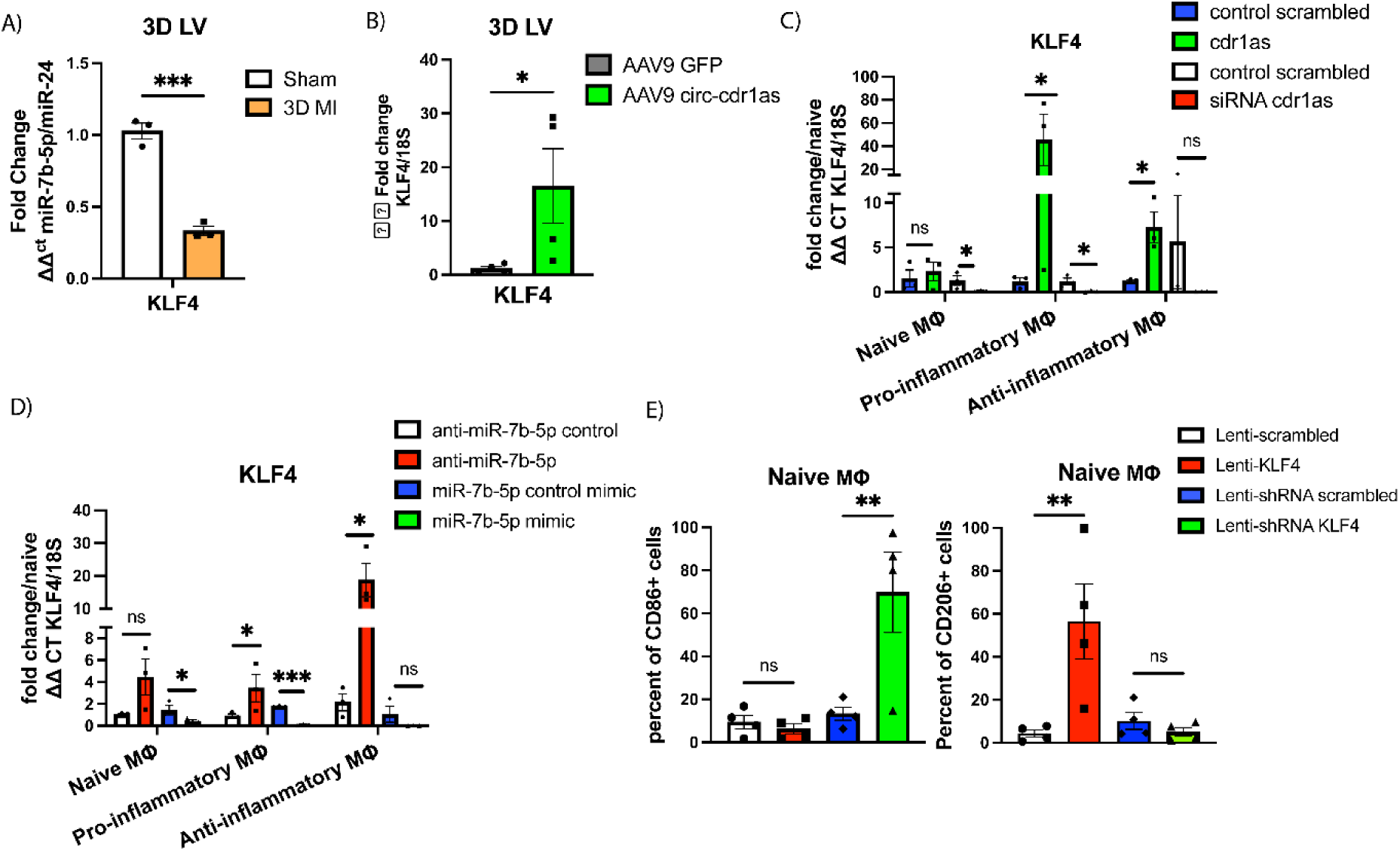
Circ-cdr1as modulates expression of miR-7b-5p target KLF4 in the heart 3D post-MI and in BMDMs. (A) qRT-qPCR of KLF4 expression in 3D post-MI left ventricular (LV) tissue compared to sham, normalized to 18S. N=3/group. (Unpaired T-test) (B) RT-qPCR of KLF4 expression in 3D post-MI left ventricular tissue in AAV9-GFP and AAV9-circ-cdr1as groups, normalized to 18S. N=4 (Unpaired-t-test). (C) Changes in KLF4 expression in naïve, pro-inflammatory, and anti-inflammatory macrophages overexpressing circ-cdr1as or knockdown of circ-cdr1as, normalized to 18S. N= 3/group (One-way ANOVA). (D) Changes in KLF4 expression in naïve, pro-inflammatory, and anti-inflammatory macrophages treated with miR-7b-5p mimic or anti-mir-7b-5p, normalized to miR-24. N= 3/group (One-way ANOVA). (E) FACS analysis of F4/80/CD86 + cells (pro-inflammatory MΦ marker) or (F) F4/80/CD206+ cells (anti-inflammatory MΦ marker) in naïve macrophages treated with lentivirus KFL4 or shRNA KLF4 and their respective controls. N=3-4/group (One-way ANOVA). Data are mean ± SEM. NS, non-significant, *p<0.05, ** p<0.01, *** p<0.001. MΦ, macrophages; pro-inflammatory marker: CD86; anti-inflammatory marker: CD206.

**Figure 8.**
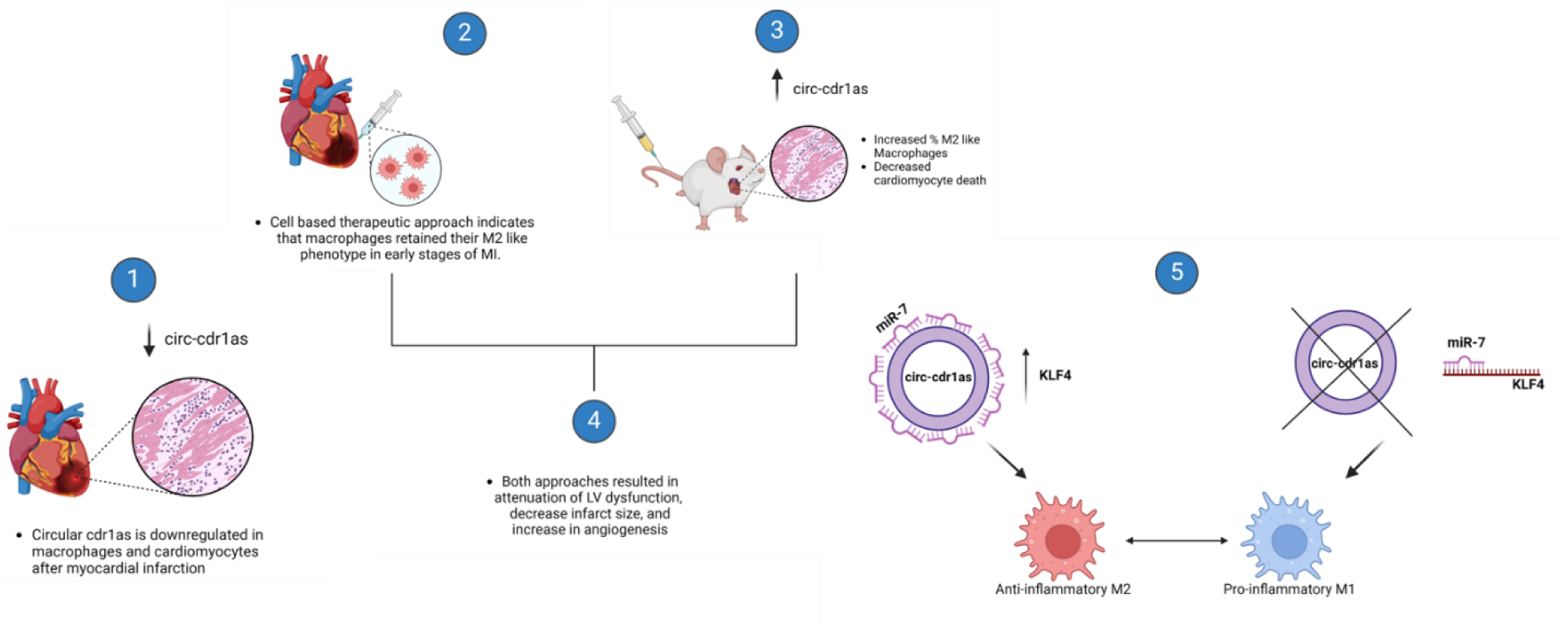
Graphical Summary.

## Discussion

Macrophages are key cellular elements of chronic inflammatory responses associated with cardiovascular disease, cancer, and other inflammatory conditions ^6,8,56,57^. These cells exhibit both pro-inflammatory (M1) and anti-inflammatory (M2) phenotypes, which are essential for tissue repair and immune regulation. However, sustained activation of pro-inflammatory macrophages can lead to adverse tissue remodeling that can ultimately contribute to deterioration of cardiac function and heart failure ^4–6^. Despite their importance, the molecular mechanisms that govern macrophage polarization, particularly in the context of cardiac injury, remain not fully understood.

Emerging evidence indicates that non-coding RNAs (ncRNAs), such as microRNAs (miRNAs) and long non-coding RNAs (lncRNAs), play pivotal roles in modulating macrophage polarization. Most studies have focused on the role of miRNAs and lncRNAs in shaping macrophage polarization and plasticity ^58–61^. In particular, a novel class of ncRNAs, circular RNAs, have garnered attention due to their unique structural properties and differential expression in cardiovascular disease, cancer, and inflammatory diseases ^14,19,62,63^. Recent studies have highlighted circRNAs’ potential to modulate macrophage polarization, suggesting that they might influence immune responses in cardiovascular disease ^28,64–66^. Reportedly, circRNA-RNF19B expression in M1 macrophages was significantly higher than that in M2 macrophages. Interestingly, expression increased when M2 macrophages were polarized to an M1 phenotype. Knockdown of circRNF19B following M1 activation downregulated expression of M1 macrophages markers and elevated the expression of M2 macrophages markers ^64^. Another study found that circular antisense non-coding RNA in the INK4 locus (circANRIL) confers atheroprotection by regulating ribosomal biogenesis in macrophages. CircANRIL binds to pescadillo homologue 1 (PES1), an essential 60S-preribosomal assembly factor to prevent exonuclease-mediated pre-rRNA processing and ribosome biogenesis ^66^. Additionally, we previously reported for the first time the role of circ-cdr1as in modulation of macrophage phenotype and overexpression of circ-cdr1as promotes phenotypic switching to an anti-inflammatory phenotype in BMDMs ^28^. However, the role of circ-cdr1as in modulation of macrophage function during cardiac injury has yet to be established.

In acute MI, circ-cdr1as was demonstrated to increase cardiac infarct size and apoptosis 24 hours after cardiac injury by LAD ligation. Further in vitro studies on mouse cardiac myocyte cell line supported circ-cdr1as as pro-apoptotic by preventing the inhibition of pro-apoptotic targets of miR-7, poly ADP ribose polymerase (PARP) and member of the SP/KLF family of transcription factors SP1^50^. Although this study did reveal a possible role of circ-cdr1as/miR-7 in cardiomyocytes during acute MI, it is important to note that these observations were limited to the first initial 24 hours of cardiac injury. Another study between circ-cdr1as and post-MI cardiac function in pigs demonstrated a negative correlation between circ-cdr1as and infarct size. They observed a downregulation of circ-cdr1as 7 days post-MI ^67^. These findings support an integral role of circ-cdr1as in cardiac dysfunction. However, mechanistic underpinning and leveraging circ-cdr1as to modulate cardiac inflammation and improve post-MI function is not established, and our studies provide direct evidence on these aspects of cdr1as function.

The CDR1 (cerebellar degeneration related 1) gene is associated with cerebellar degeneration, Alzheimer’s diseases, and hepatocellular carcinoma. The CDR1 gene is in the X chromosome and is highly conserved between humans and mice. The CDR1 protein is elevated in leukocytes in Alzheimer’s patients ^68^. Circ-cdr1as is a closed circular RNA formed from the antisense transcript of the CDR1 gene and our previous work indicated that expression levels of circular form of circ-cdr1as were changed during macrophage polarization without affecting the linear transcript ^28^. Bioinformatic analysis of the functional role of circ-cdr1as in cancer suggests that this circRNA plays a role in immune and stromal cell infiltration within the tumor tissue. Tissues with high circ-CDR1as expression had a higher ratio of anti-inflammatory macrophages and circ-CDR1as expression was negatively correlated with CD8^+^ T cells ^49^.

In the current study, our findings suggest that circ-cdr1as may play a central role in modulating the inflammatory environment after MI. We demonstrate that circ-cdr1as is significantly repressed during the early phase of MI, particularly in cardiomyocytes and macrophages. at 3 days post-MI, which aligns with the peak of inflammation. This temporal decrease in circ-cdr1as expression occurred during the phase dominated by pro-inflammatory macrophages. Interestingly, expression of circ-cdr1as increased in macrophages after day 7, corresponding with the start of resolution of inflammation.

Anti-inflammatory MΦs secrete high levels of anti-inflammatory cytokines and growth factors including VEGF, known to promote angiogenesis ^8,69^. Anti-inflammatory MΦs can also produce myeloid-derived growth factor know to protect cardiomyocytes from cell death ^45^, and secrete TGFβ1 leading to upregulation of α-SMA to aid in fibroblast differentiation and facilitate tissue repair ^70^. We observed that myocardial injection of macrophages overexpressing circ-cdr1as maintained their anti-inflammatory phenotype in inflammatory conditions following cardiac injury. Myocardial injection of circ-cdr1as MΦs enhanced angiogenesis, reduce infarct size, and attenuate LV dysfunction after MI. Additionally, our findings indicate that overall cardiac overexpression of circ-cdr1as improved left ventricular function and reduced infarct size at 21- and 28-days post-MI. There was also increased myocardial neovascularization, supporting the idea that circ-cdr1as may promote tissue repair by enhancing angiogenesis. Interestingly, we observed an increase in the number of anti-inflammatory macrophages at the injury site, suggesting that circ-cdr1as may influence the inflammatory environment within the heart. This could be the result of direct overexpression in cardiac cells or cross talk between cardiac cells to aid in the cardioprotective effects of circ-cdr1as. Notably, cardiomyocytes are known to release factors that promote phenotypic switching in macrophages, differentiation in fibroblast, and stimulate angiogenesis ^8,71,72^. However, more research is needed to understand the relationship between them.

The most established function of circRNAs located in the cytoplasm is acting as miRNAs sponges ^73–76^. Therefore, we investigated the potential relationship between circ-cdr1as and miRNAs and identified miR-7 as a target. Previous work has shown circ-cdr1as has over 70 binding sites and their interaction is highly conserved between species ^49,51,77^. Recent studies demonstrated that miR-7 is a key regulator of the immune response influencing T cell activation, macrophage function, and dendritic cell maturation in inflammation-related diseases ^78–82^. However, the role of miR-7 in modulation of the inflammatory response in cardiovascular disease is largely unexplored. In the current study, we demonstrate that circ-cdr1as significantly represses miR-7 expression in macrophages and cardiomyocytes following MI and in BMDMs. We also identified that treatment with miR-7 mimic in BMDMs stimulated these cells to phenotypically switch to a more pro-inflammatory phenotype; meanwhile, treatment with anti-miR-7 has the opposite effect. Furthermore, inhibition of miR-7 upregulates Krüppel-like factor 4 (Klf4), a key transcription factor involved in immune cell regulation and inflammation. We show that circ-cdr1as overexpression results in the upregulation of Klf4 in naïve, pro-, and anti-inflammatory macrophages, suggesting a mechanistic link between circ-cdr1as, miR-7, and Klf4 in regulating macrophage phenotype during cardiac injury. Klf4 is an evolutionarily conserved zinc finger-containing transcription factor that regulates cell growth, proliferation, and differentiation ^83^. Of note, miR-7 deficiency and Klf4 upregulation were associated with changes in immune cell composition and increase proportion of MHC anti-inflammatory macrophages in an acute lung injury model. Studies have previously reported that Klf4 is a critical regulator in LPS-induced inflammatory response ^84^, regulate the transduction of NF-κB pathway ^85^, and may acts as a transcriptional regulator to upregulate TGFβ1 and IL-10 ^86,87^. This study reports that Klf4 is downregulated in LV tissue 3 days post-MI and upregulated in mice with cardiac overexpression of circ-cdr1as. Moreover, we observe that the upregulation of Klf4 in macrophages following miR-7 inhibition is associated with an increased percentage of CD206+ anti-inflammatory macrophages and a decrease in CD86+ pro-inflammatory cells. These findings are consistent with the idea that Klf4 promotes anti-inflammatory macrophage polarization and supports tissue repair in response to injury. Additionally, the transcriptional upregulation of anti-inflammatory markers (Arg-1 and CD206) and the downregulation of pro-inflammatory markers (Ly6C and INOS) further support the notion that Klf4 plays a central role in macrophage polarization during cardiac injury. Pharmacological inhibition of klf4 using kenpaullone, a small molecule known to inhibit CDK1 and GSK-3β, which has been shown to modulate Klf4 expression ^53–55^, provided similar readouts, providing further evidence for the central role of Klf4 in regulating macrophage phenotypic switching. This aligns with previous research showing that Klf4 is a critical regulator of inflammation and immune responses, particularly in the context of lung injury and sepsis ^86,87,86,87^. Furthermore, modulation of circ-cdr1as levels in our study reveals a crucial role of circ-cdr1as/miR-7/Klf4 in cardiac injury and macrophage phenotype.

While our study provides significant insights into the role of circ-cdr1as in macrophage polarization and cardiac injury, some limitations must be acknowledged. First, we cannot definitively attribute all observed changes in macrophage polarization to miR-7 and Klf4 alone, as circ-cdr1as may also interact with other miRNAs or molecules that influence macrophage behavior. Further studies using transcriptomic or proteomic approaches could help identify additional downstream targets of circ-cdr1as. Additionally, although we have shown the effects of circ-cdr1as overexpression in BMDMs and macrophages during MI, it remains unclear how long these macrophages maintain their phenotype post-injury and whether these effects translate into long-term functional improvements in the heart. Longitudinal studies investigating the persistence of macrophage phenotype and the impact on cardiac remodeling long term would be valuable. Another important consideration is the role of Klf4 in the resolution of inflammation and tissue repair in the heart. Although our findings suggest that Klf4 promotes an anti-inflammatory macrophage phenotype, more detailed studies are needed to explore the specific signaling pathways through which Klf4 modulates macrophage function in cardiac injury. Furthermore, the potential therapeutic implications of targeting the circ-cdr1as/miR-7/Klf4 axis in promoting cardiac repair and reducing infarct size should be explored.

## Conclusion

In summary, our findings shed light on the regulatory roles of circRNAs, particularly circ-cdr1as, as a possible regulator of macrophage function in the context of cardiac injury, potentially influencing the resolution of inflammation and tissue repair. This study suggests that overexpression of circ-cdr1as may be cardioprotective by reducing cardiomyocyte apoptosis, enhancing angiogenesis, limiting infarct size, increasing percentage of anti-inflammatory macrophages, and overall preserving post-MI cardiac function. We present evidence that circ-cdr1as modulates macrophage phenotypic switching through the miR-7/Klf4 axis suggesting that targeting this pathway could provide therapeutic benefits in cardiovascular diseases.

## Author Contributions

C. Gonzalez and R. Kishore conceived and designed the project. M. Cimini assisted in the experiments, data interpretation and analysis. C. Thej, M. Truongcao, C. Benedict, A Rai assisted in the experiments. Z. Cheng, V.N.S. Garikipati and D. Joladarashi assisted in data interpretation. C. Gonzalez and R. Kishore wrote and edited the article.

