## Supplemental figures and tables for "Circular RNA Circ-Cdr1as modulates Macrophage phenotype and Cardiac Reparative Function by Circ-Cdr1as-miR-7-Klf4 pathway"

### SUPPLEMENTAL MATERIAL

### 1. Expanded Materials and Methods

### 2. Supplemental Tables

### 3. Supplemental Figures with Figure Legends

### 3. Major Resources

### EXPANDED METHODS

#### Myocardial infarction surgery and treatments:

Animals are acclimated to the house environment for at least a week prior to surgery. Mice were anesthetized with 2% isoflurane and orally intubated with an intubated with a 22 G i.v. catheter and artificially ventilated with a respirator (Harvard Apparatus). To provide analgesia, buprenorphine (0.5 mg/kg) was injected subcutaneously before the operation. A left intercostal thoracotomy was performed, and the ribs were retracted with 5-0 polypropylene sutures to open the chest. After the pericardium was opened, the left anterior descending artery (LAD) was ligated distal to the bifurcation between the LAD and diagonal branch using 8-0 polypropylene sutures through a dissecting microscope. After positive end-expiratory pressure was applied to inflate the lung fully, the chest was closed with 7-0 polypropylene sutures. A 22 G syringe was used to evacuate air from the chest cavity. The mice in the sham group underwent the same procedure except for the LAD ligation.

For the macrophage cell therapy study, immediately after LAD ligation, each mouse received intramyocardial injection of ~500,000 AAV2 circ-cdr1as macrophages (N=17) or AAV2 vehicle macrophages (N=17) or saline (N=17) in a total volume of 25 µl at 5 different sites (basal anterior, mid anterior, mid lateral, apical anterior, and apical lateral).

For the second *in vivo* study, mice received $1x10^{11}$vp/mL of AAV9 circ-cdr1as (N=15) or AAV9 vehicle (N=15) by tail vein injection in a total volume of 200 µL14 days prior to MI as previously described [31-34]. Briefly, prior to injection animals were warmed for 5-10 minutes to dilate the veins by placing a heating pad under the cage. Then each mouse was placed in a restraint device and injected slowly with either AAV9 circ-cdr1as or AAV9 vehicle and the device was cleaned between each mouse. Following injection, we applied a firm compression on the injection site with a clean gauze to minimize bleeding and then returned them to their cage. The mice were then monitored until they resumed normal activity.

The survival rate of MI surgery was 83.6%. At the endpoint, body weight, heart tissue weight, and tibia length were measured and heart tissues were collected under anesthesia with Avertin® (222-Tribromoethanol; T48402; MilliporeSigma, USA). The animals were then euthanized following the procedure from our approved IACUC protocol.

***Circ-cdr1as Expression Plasmid and AAV Construction***

Our circ-cdr1as expression plasmid was generated as previously described [28]. Briefly, we used pcDNA3.1 (+) Lacasse2 MCS exon vector (Addgene plasmid # 69893; http://n2t.net/addgene:69893; RRID:Addgene_69893) [35] with fragment insertion in *PacI* and *SacII* sites of pcDNA3.1 (+) Lacasse2 MCS exon vector (Figure S2A). For *in vivo* expression, the sequence was cloned into a self-complementary AAV backbone plasmid (pTRUFR) and expression of the circRNA is driven by a CMV promoter and flanked by AAV9 ITRs (*in vivo* tail vein injections) or AAV2 (*ex vivo* overexpression in macrophages). AAV9-GFP and AAV2-empty vector were used as controls. All viruses were produced using the triple transfection technique [36]. In brief, HEK293 cells were plated at $1 \times 10^7$ cells/15 cm plate one day prior to transfection and polyethyleneimine (Linear PEI, MW 25 kDa, Sigma-Aldrich) was used as the transfection reagent. For each 15 cm plate, the following reagents were added: 12 μg XX6-80 [37], 10 μg pXR9 [38], and 6 μg pTRUFR WT-circ-cdr1as. These plasmids were combined in 500 μl of DMEM without serum or antibiotics and 100 μl of PEI reagent (1 mg/ml pH5.0) was added, mixed, and set aside for 5–10 min. Without changing the media on the 15 cm plates, the DNA-PEI reagent was added to the cells drop-wise. Cells were then harvested 48–72 h after transfection, collected by centrifugation and the cell pellets were re-suspended in water and freeze-thawed once prior to sonication (Branson Sonifier 250, VWR Scientific). DNase I (Sigma, MO) and MgCl2 were then added before incubating at 37 °C for 60 min. The viruses were purified by Iodixanol step gradient centrifugation5 followed by FPLC using HiTrap Q HP anion exchange chromatography column and AKTA Pure FPLC system (GE Healthcare Life Sciences). The purified virus particles were recovered and filtered through a 0.22 μM filter and quantified by OD 260. The viruses were aliquoted and stored at −80 °C in phosphate buffered saline (PBS) containing 5% sorbitol. AAV titers were determined by real-time PCR on plasmid genomes using the SYBR Green Master Mix (Roche).

#### *Left Ventricle Heart Tissue Collection and Cardiac Cell Isolations*

To isolate RNA from the LV tissue, the heart was quickly removed from the chest and perfused with phosphate buffered saline (PBS) 3x. The LV tissue was excised and minced prior to RNA extraction. Adult mouse cardiomyocytes, endothelial cells, and fibroblast were isolated as previously described [39]. Once cardiomyocytes are isolated, non-myocyte supernatant fractions of LV digestion was centrifuged (300g for 5 mins), and cardiac endothelial cells were isolated by magnetic bead separation using CD31+ beads. Afterwards, fibroblasts were isolated according to the previously described protocol. To obtain macrophages from the heart, the LV was excised, weighed, and minced with a fine scissor prior to digestion with collagenase type 2 (250 U/mL; Worthington), collagenase type XI (125 U/mL; Worthington), deoxyribonuclease I (60 U/mL Worthington), and hyaluronidase (60 U/mL;) for 10 minutes at 37 °C. The tissue was then allowed to settle to the bottom, the supernatant was collected and passed through a 40μm mesh filter into a 50mL conical tube. Then 2 ml of stopping buffer (10% FBS in PBS) was added to the collected supernatant. The 15 mL tube with the settled tissue then went through another round of digestion. This process was repeated 4 times and then the collected supernatant was centrifuged at 50G for 5 minutes. The supernatant was then collected into another 50 mL conical tube and centrifuged at 500 g for 10 mins. The supernatant was discarded, and the pellet was resuspended in 1 mL of ACK lysis buffer (Gibco) for 5 mins. Then, 5mL of HEPES stock solution (Gibco) was added and centrifuged for 400 g for 5 mins at 4 °C. The cells were then prepared for FACS.

#### *FACS Analysis of Cardiac and Bone Marrow Derived Macrophages*

After isolation of macrophages from the heart as described above, the cells were pre-incubated with 0.25μg of TrueStain FcX PLUS (anti-mouse CD16/32) antibody (Biolegend) in 100 μL volume for 10 minutes on ice. The cells were washed with staining buffer and incubated with F4/80 antibody at 4ºC for 30 minutes then subsequently washed with staining buffer 3x and resuspended in 1 mL of staining buffer. The cells were then sorted using BD FACS ARIA IIμ based on F4/80+ marker. The sorted cells were then collected for RNA extraction.

Flow cytometry was used to characterize and identify the percentage of GFP+, F4/80+, CD206+, CD86+ or Ly6c+ cells in cardiac macrophages and bone marrow derived macrophages. Briefly, after isolation of macrophages from the heart as described above or harvested BMDMs, the cells counted and $1.5x10^6$ cells were collected for analysis using BD FACSymphony A5 Cell Analyzer and FACSDiva (BD Biosciences, USA). The cells were pre-incubated with 0.25μg of TrueStain FcX PLUS (anti-mouse CD16/32) antibody (Biolegend) in 100 μL volume for 10 minutes on ice. The cells were washed with staining buffer and incubated with the following antibodies at 4ºC for 30 minutes then subsequently washed with staining buffer 3x and resuspended in 1 mL of staining buffer. The samples were analyzed using the BD FACS AriaTM (BD Biosciences, USA). Appropriate compensation controls were used to negate spectral overlap and gating strategies were based on fluorescence minus one (FMO).

***Tissue preparation and Immunohistochemistry***

Mouse heart tissue samples were fixed in 4% paraformaldehyde (PFA) for at least 48 hours. The tissue was washed with PBS 3x and then placed in 10% sucrose for one hour, washed 3x, then placed in 20% sucrose for hour, washed 3x, and placed in 32% sucrose O/N. The tissue was then dissected and embedded with OCT in a labeled cryomold. The cryomold was frozen in liquid nitrogen then temporarily stored in dry ice and wrapped in foil for long term storage in -80ºC. Cardiac tissues were cross-sectioned into 4–5 μm-thick slides.

Masson Trichrome staining (Sigma Aldrich, USA) was performed following the manufacturer's instructions and percent of fibrotic area was determined by fibrosis area/total area using Image J software. Masson Trichrome staining images were acquired with Nikon stereomicroscope at 1X (scar quantification).

For identification of new capillary network, sections were stained with CD31 and α-SMA. For identification of immune cells and anti-inflammatory macrophages, sections were heat-induced for antigen retrieval prior to immunolabeling for CD45 (immune cells), CD163, CD68, and CD206 (anti-inflammatory macrophage markers) were used. For identification of GFP+ cells, sections were stained with GFP antibody. To determine AAV9 GFP+ cardiomyocytes, sections were stained with GFP in green and Troponin T in red. To determine cell death in cardiomyocytes, sections were stained with TUNEL in red (Roche) and Troponin T in green. To determine cardiomyocyte hypertrophy, sections were stained with wheat germ agglutinin (WGA). Nuclei were counter-stained with 4,6-diamidino-2-phenylindole (DAPI, 1:10,000, Sigma Aldrich) then mounted using VECTASHIELD® PLUS Antifade Mounting Medium (Vector Laboratories, CA, USA). Images were acquired using 4X, 10X, 20X, and 40X using EVOS 7000 (Thermo Fisher Scientific). Quantitative image analysis was performed with NIH ImageJ by scoring random multiple imaging fields.

***Klf4 inhibition with Kenpaullone***

Kenpaullone (MCE) is a potent inhibitor of CDK1/cyclin B and GSK-3β that has been shown to inhibit klf4. We treated BMDMs with 100nM of kenpaullone for 24 hours prior to BMDMs polarization. Cells were harvested and changes in the expression or transcription of macrophage markers were measured by FACS or RT-qPCR, respectively.

***RNAseR Treatment, Reverse transcription, and RT-qPCR***

Total RNA was isolated from cells using miRNeasy Mini Kit (Qiagen). NanoDrop-1000 (Thermo Scientific, USA) was used to identify RNA concentration and purity determined by the 260 A/280 A ratio. For RNase R treatment, 500 ng was incubated 30 mins at 37°C with or without 2.5 U of Rnase R (Epicentre Technologies). High-capacity cDNA Reverse Transcription Kit (Applied Biosystems) was used to obtain cDNA and quantitative PCR (qPCR) was performed on StepOnePLus Real-Time PCR system (Applied Biosystems 7000 apparatus) using the Fast SYBR™ Green Master Mix (Applied ThermoFisher, USA) according to the manufacturer's instructions. To quantify expression of circRNA transcripts, divergent primers were designed to amplify across the backsplicing junction. To quantify linear transcripts, convergent primers were designed to amplify exonic sequences that are not present in the circular RNA. Expression was quantified using fold change of $2^{e\ (delta\ ct)}$ compared to control, normalized to 18S rRNA used as a reference gene.

For detection of miRNAs, TaqMan™ Advanced miRNA cDNA Synthesis Kit was used (Thermofisher Cat). Quantitative PCR (qPCR) was performed on StepOnePLus Real-Time PCR system (Applied Biosystems 7000 apparatus) using the TaqMan™ Fast Advanced Master Mix for qPCR (Thermofisher) according to the manufacturer's instructions. To detect miR-7b-3p expression primer mmu-miR-7b-5p (Thermofisher) was used and expression was quantified using fold change of $2^{e\ (delta\ ct)}$ compared to control, normalized to miR-24-3p (Assay ID mmu481011_mir, Thermofisher) used as a reference gene.

### SUPPLEMENTARY TABLES

**Supplementary Table 1:** The echocardiographic parameters measured in mice at baseline for sham (N=5), saline (N=7), AAV2 vehicle (N=10), and AAV2 circ-cdr1as groups (N=10).

| Baseline | Sham | Saline | AAV2 vehicle | AAV2 circ-cdr1as | Sham | Saline | AAV2 vehicle | AAV2 circ-cdr1as |
|---|---|---|---|---|---|---|---|---|
| ***Parameters*** | **Mean** | | | | **SEM** | | | |
| ***EF (%)*** | 59.3 | 60.0 | 59.5 | 58.0 | 1.1 | 4.3 | 1.7 | 1.4 |
| ***FS (%)*** | 31.0 | 31.3 | 30.7 | 29.9 | 0.7 | 2.8 | 1.1 | 0.7 |
| ***CO (mL/min)*** | 18.2 | 16.0 | 13.6 | 16.4 | 1.9 | 4.2 | 2.0 | 3.6 |
| ***LVID;d (mm)*** | 3.9 | 3.7 | 3.3 | 3.6 | 0.1 | 0.4 | 0.2 | 0.3 |
| ***LVID;s (mm)*** | 2.7 | 2.5 | 2.3 | 2.7 | 0.1 | 0.4 | 0.2 | 0.3 |

**Supplementary Table 2:** The echocardiographic parameters measured in mice at 7 days post MI for sham (N=5), saline (N=7), AAV2 vehicle (N=10), and AAV2 circ-cdr1as groups (N=10).

| 7D | Sham | Saline | AAV2 vehicle | AAV2 circ-cdr1as | Sham | Saline | AAV2 vehicle | AAV2 circ-cdr1as |
|---|---|---|---|---|---|---|---|---|
| ***Parameters*** | **Mean** | | | | **SEM** | | | |
| ***EF (%)*** | 57.7 | 29.8 | 30.5 | 28.8 | 0.9 | 6.2 | 2.5 | 4.3 |
| ***FS (%)*** | 29.7 | 14.0 | 14.0 | 13.4 | 0.5 | 3.2 | 1.1 | 2.0 |
| ***CO (mL/min)*** | 15.7 | 14.0 | 10.1 | 13.0 | 2.5 | 2.7 | 3.3 | 2.5 |
| ***LVID;d (mm)*** | 3.5 | 4.5 | 3.9 | 4.6 | 0.2 | 0.4 | 0.6 | 0.7 |
| ***LVID;s (mm)*** | 2.5 | 3.9 | 3.3 | 4.0 | 0.2 | 0.5 | 0.5 | 0.7 |

**Supplementary Table 3:** The echocardiographic parameters measured in mice at 21 days post MI for sham (N=5), saline (N=7), AAV2 vehicle (N=10), and AAV2 circ-cdr1as groups (N=10).

| 21D | Sham | Saline | AAV2 vehicle | AAV2 circ-cdr1as | Sham | Saline | AAV2 vehicle | AAV2 circ-cdr1as |
|---|---|---|---|---|---|---|---|---|
| ***Parameters*** | **Mean** | | | | **SEM** | | | |
| ***EF (%)*** | 56.5 | 23.4 | 27.3 | 39.9 | 1.0 | 6.5 | 5.9 | 4.1 |
| ***FS (%)*** | 29.0 | 10.7 | 12.7 | 19.1 | 0.7 | 3.2 | 2.8 | 2.1 |
| ***CO (mL/min)*** | 17.1 | 11.2 | 13.9 | 13.8 | 2.6 | 4.7 | 4.0 | 1.1 |
| ***LVID;d (mm)*** | 3.7 | 4.5 | 4.7 | 3.9 | 0.3 | 0.9 | 1.1 | 0.3 |
| ***LVID;s (mm)*** | 2.6 | 4.1 | 4.1 | 3.1 | 0.2 | 0.9 | 1.1 | 0.3 |

**Supplementary Table 4:** The echocardiographic parameters measured in mice at 28 days post MI for sham (N=5), saline (N=7), AAV2 vehicle (N=10), and AAV2 circ-cdr1as groups (N=10).

| **28D** | Sham | Saline | AAV2 vehicle | AAV2 circ-cdr1as | Sham | Saline | AAV2 vehicle | AAV2 circ-cdr1as |
|---|---|---|---|---|---|---|---|---|
| ***Parameters*** | **Mean** | | | | **SEM** | | | |
| ***EF (%)*** | 54.7 | 19.3 | 28.0 | 43.3 | 1.3 | 6.3 | 6.8 | 5.7 |
| ***FS (%)*** | 27.9 | 8.8 | 13.0 | 21.2 | 0.8 | 3.0 | 3.4 | 3.1 |
| ***CO (mL/min)*** | 16.6 | 10.3 | 13.9 | 18.2 | 0.8 | 1.4 | 7.5 | 4.6 |
| ***LVID;d (mm)*** | 3.8 | 4.8 | 4.5 | 4.2 | 0.1 | 0.6 | 1.1 | 0.5 |
| ***LVID;s (mm)*** | 2.8 | 4.4 | 3.9 | 3.3 | 0.1 | 0.7 | 1.0 | 0.5 |

**Supplementary Table 5:** The echocardiographic parameters measured in mice at baseline for saline (N=5), AAV9 GFP (N=7), and AAV9 circ-cdr1as groups (N=7).

| **Baseline** | Saline | AAV9 GFP | AAV9 circ-cdr1as | Saline | AAV9 GFP | AAV9 circ-cdr1as |
|---|---|---|---|---|---|---|
| ***Parameters*** | **Mean** | | | **SEM** | | |
| ***EF (%)*** | 62.1 | 60.0 | 61.0 | 5.6 | 2.4 | 2.0 |
| ***FS (%)*** | 32.9 | 31.3 | 31.1 | 3.8 | 1.4 | 2.8 |
| ***CO (mL/min)*** | 16.6 | 17.6 | 16.0 | 4.3 | 2.3 | 4.6 |
| ***LVID;d (mm)*** | 3.7 | 4.0 | 3.7 | 0.5 | 0.1 | 0.3 |
| ***LVID;s (mm)*** | 2.5 | 2.7 | 2.5 | 0.5 | 0.1 | 0.2 |

**Supplementary Table 6:** The echocardiographic parameters measured in mice at 7 days post MI for saline (N=5), AAV9 GFP (N=7), and AAV9 circ-cdr1as groups (N=7).

| **7D** | Saline | AAV9 GFP | AAV9 circ-cdr1as | Saline | AAV9 GFP | AAV9 circ-cdr1as |
|---|---|---|---|---|---|---|
| ***Parameters*** | **Mean** | | | **SEM** | | |
| ***EF (%)*** | 30.9 | 22.6 | 26.8 | 6.2 | 2.6 | 1.8 |
| ***FS (%)*** | 14.6 | 10.7 | 12.8 | 3.3 | 0.9 | 0.7 |
| ***CO (mL/min)*** | 18.4 | 12.3 | 15.6 | 4.4 | 5.8 | 3.6 |
| ***LVID;d (mm)*** | 4.6 | 4.7 | 4.8 | 0.4 | 0.8 | 0.7 |
| ***LVID;s (mm)*** | 3.9 | 4.0 | 4.2 | 0.4 | 0.8 | 0.7 |

**Supplementary Table 7:** The echocardiographic parameters measured in mice at 21 days post MI for saline (N=5), AAV9 GFP (N=7), and AAV9 circ-cdr1as groups (N=7).

| **21D** | Saline | AAV9 GFP | AAV9 circ-cdr1as | Saline | AAV9 GFP | AAV9 circ-cdr1as |
|---|---|---|---|---|---|---|
| ***Parameters*** | **Mean** | | | **SEM** | | |
| ***EF (%)*** | 28.5 | 17.0 | 32.8 | 7.0 | 4.1 | 4.1 |
| ***FS (%)*** | 13.5 | 8.6 | 15.3 | 3.5 | 1.6 | 2.0 |
| ***CO (mL/min)*** | 19.8 | 10.9 | 12.4 | 4.0 | 3.9 | 5.4 |
| ***LVID;d (mm)*** | 5.3 | 5.1 | 4.1 | 0.4 | 1.1 | 0.8 |
| ***LVID;s (mm)*** | 4.6 | 4.8 | 3.4 | 0.5 | 0.9 | 0.8 |

**Supplementary Table 8:** The echocardiographic parameters measured in mice at 28 days post MI for saline (N=5), AAV9 GFP (N=7), and AAV9 circ-cdr1as groups (N=7).

| **28D** | Saline | AAV9 GFP | AAV9 circ-cdr1as | Saline | AAV9 GFP | AAV9 circ-cdr1as |
|---|---|---|---|---|---|---|
| ***Parameters*** | **Mean** | | | **SEM** | | |
| ***EF (%)*** | 24.9 | 15.5 | 34.2 | 6.2 | 2.6 | 8.1 |
| ***FS (%)*** | 11.6 | 7.6 | 17.9 | 3.0 | 1.9 | 3.4 |
| ***CO (mL/min)*** | 15.2 | 11.3 | 18.0 | 4.2 | 4.2 | 6.0 |
| ***LVID;d (mm)*** | 5.0 | 5.5 | 3.9 | 0.5 | 1.2 | 0.5 |
| ***LVID;s (mm)*** | 4.5 | 5.7 | 3.3 | 0.5 | 0.9 | 0.6 |

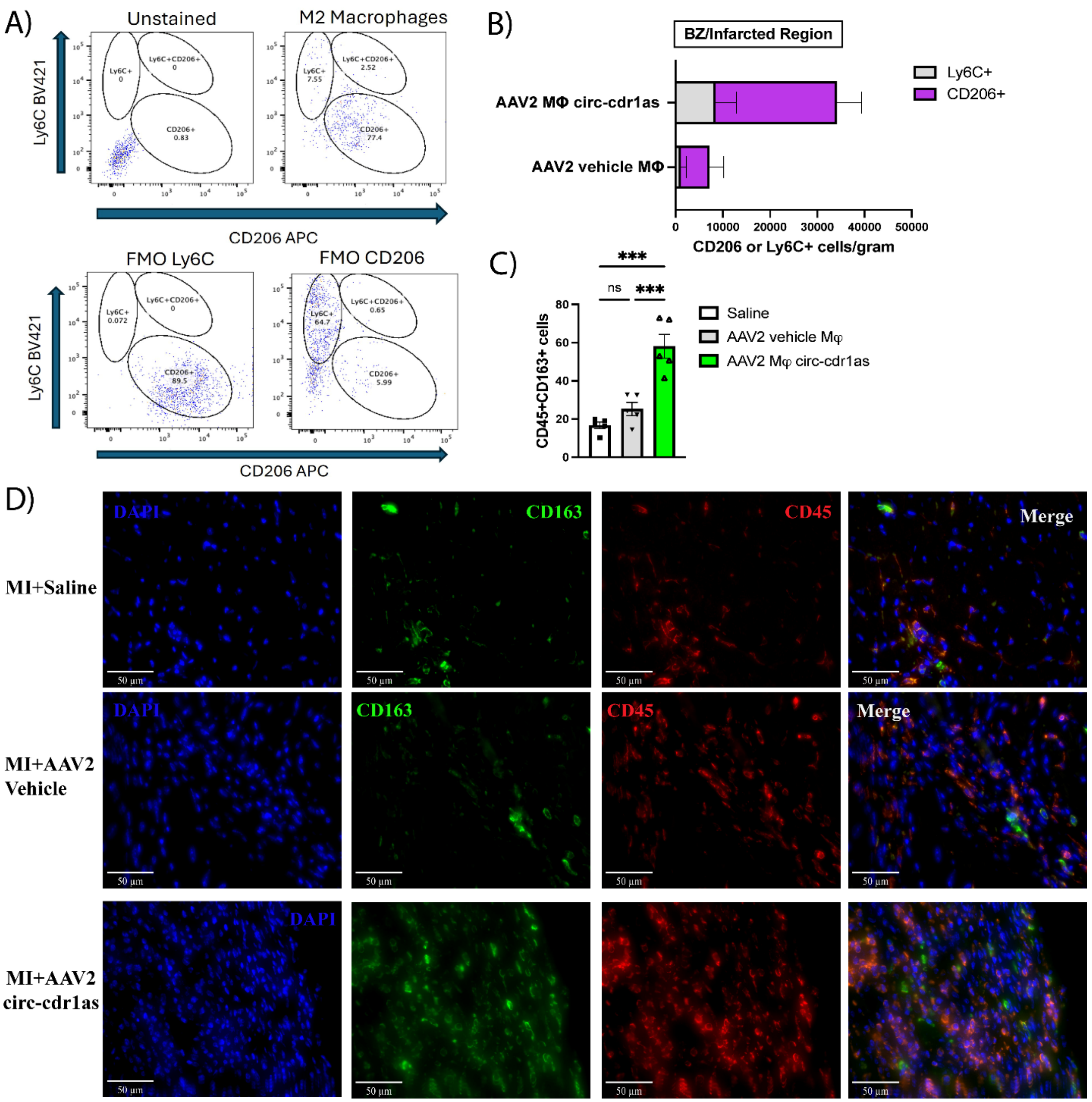


**Supplemental Figure 1 GFP+ circ-cdr1as overexpressing macrophages have an increase in CD163 macrophage marker.** (A) FACS gating strategies for Ly6c (M1 marker) and CD206 (M2) marker with fluorescent minus one (FMOs) controls. (B) FACS analysis of %GFP/CD206+ cells in the LV MI region at 5D post-MI of all normalized cells/gram for all groups. N=4/group. (C) Quantification of CD45/CD163+ cells in each group. N=5/group. (D) Representative photomicrographs (50μm) of CD163 (green) and CD45 (red) positive cells and nuclei DAPI (blue)/40X for all groups. N=5/group. One-Way ANOVA. Data are mean ± SEM. NS, non-significant, *p<0.05, ** p<0.01, *** p<0.001. Ly6C, M1 macrophage marker; CD206, M2 macrophage marker; CD45, leukocyte marker; CD163, M2 macrophage marker.

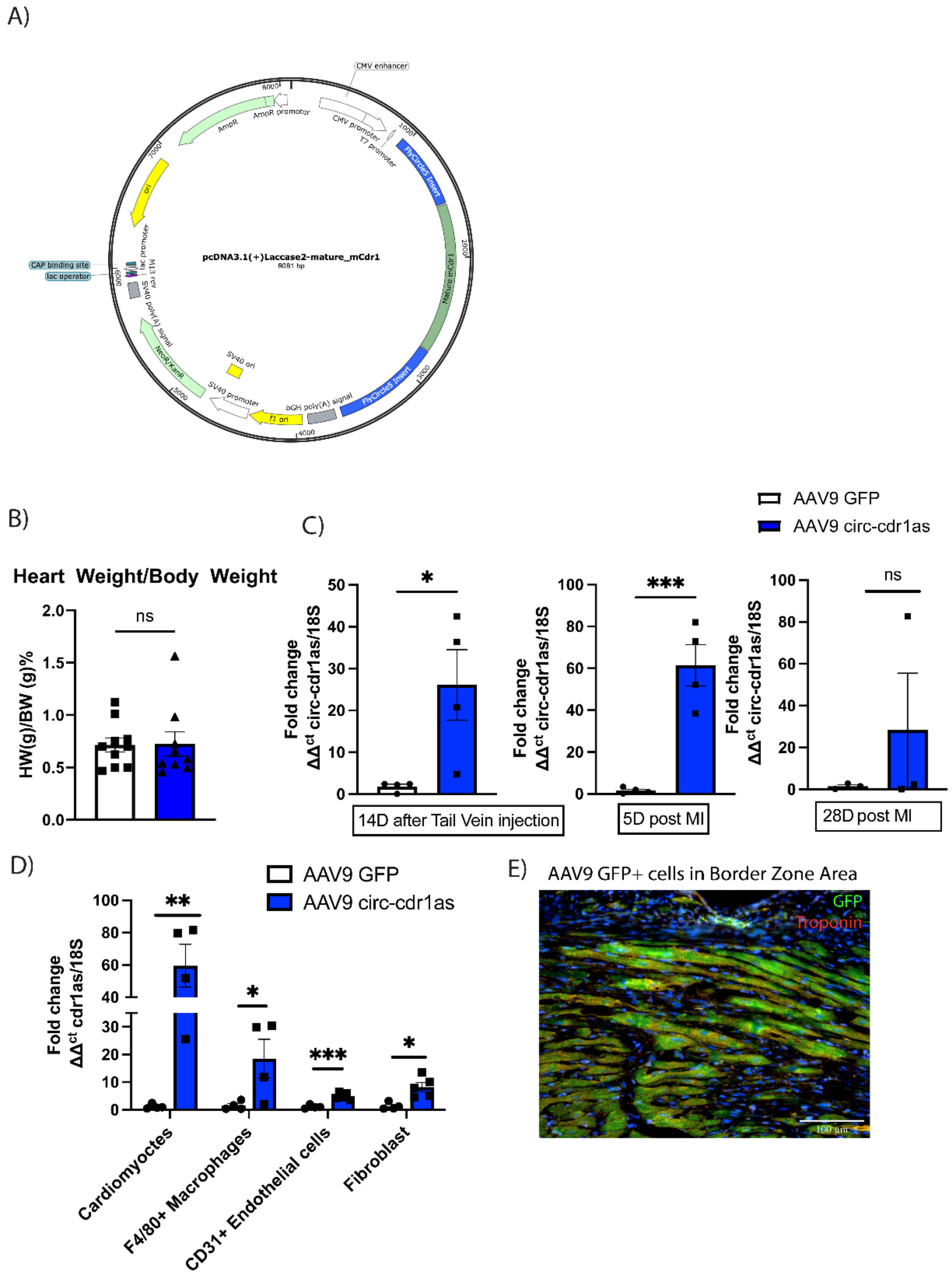


**Supplemental Figure 2 Validation of circ-cdr1as overexpression in the heart.** (A) Plasmid map of pcDNA3.1 (+) lacasse2 MCS exon vector with mature cdr1as sequence. (B) Heart and body weight ratio of AAV9 GFP or AAV9 circ-cdr1as treated mice. (C) qRT-PCR analysis of circ-cdr1as expression in AAV9 -GFP or AAV9 circ-cdr1as treated mice 14 days after tail vein injection and 5 days and 28 days (p-value 0.13) after myocardial infarction, normalized to 18S. N=3-4/group. (D) qRT-qPCR analysis of circ-cdr1as expression in cardiac cells, cardiomyocytes, macrophages (Aria sorted F4/80+ cells), endothelial cells (CD31+), and fibroblast), isolated from LV-tissue 5 days post MI from AAV9-GFP or AAV9 circ-cdr1as treated mice, normalized to 18S. N= 4-5/group. Unpaired t-test. Data are mean ± SEM. NS, non-significant, *p<0.05, ** p<0.01, *** p<0.001. (E) Representative image of GFP (green)/Troponin T (red) positive cells in border zone area in mice treated with AAV9-GFP.

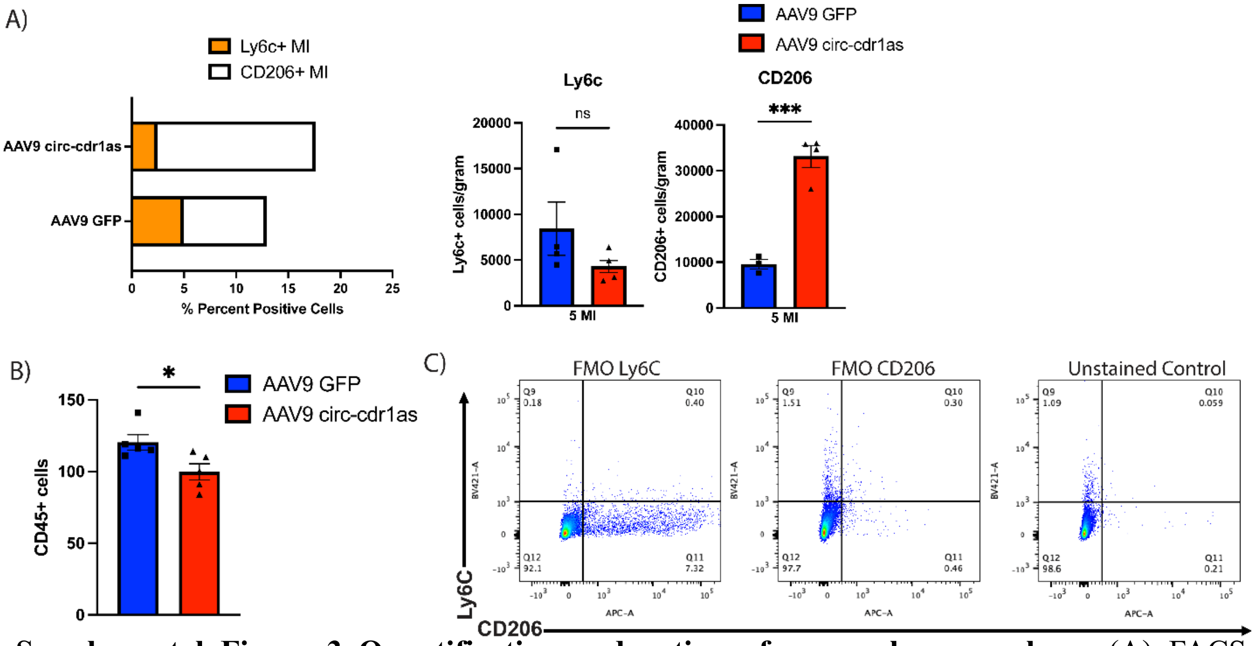


**Supplemental Figure 3 Quantification and gating of macrophage markers.** (A) FACS analysis of Ly6c+ (M1 MΦ marker) and CD206 (M2 MΦ marker) cells isolated from LV tissue 5 days post-MI and normalized cells/gram (right) for all groups. N=3-5/group (Unpaired t-test). (B) Quantification of CD45+ cells (A) N= 5/group. Unpaired t-test. (C) Fluorescent minus one (FMOs) controls for selection of gating for Ly6C and CD206+ cells. (C) Data are mean ± SEM. Unpaired t-test. NS, non-significant, NS, non-significant, *p<0.05, ** p<0.01, *** p<0.001. Ly6C, M1 macrophage marker; CD206, M2 macrophage marker; CD45, leukocyte marker.

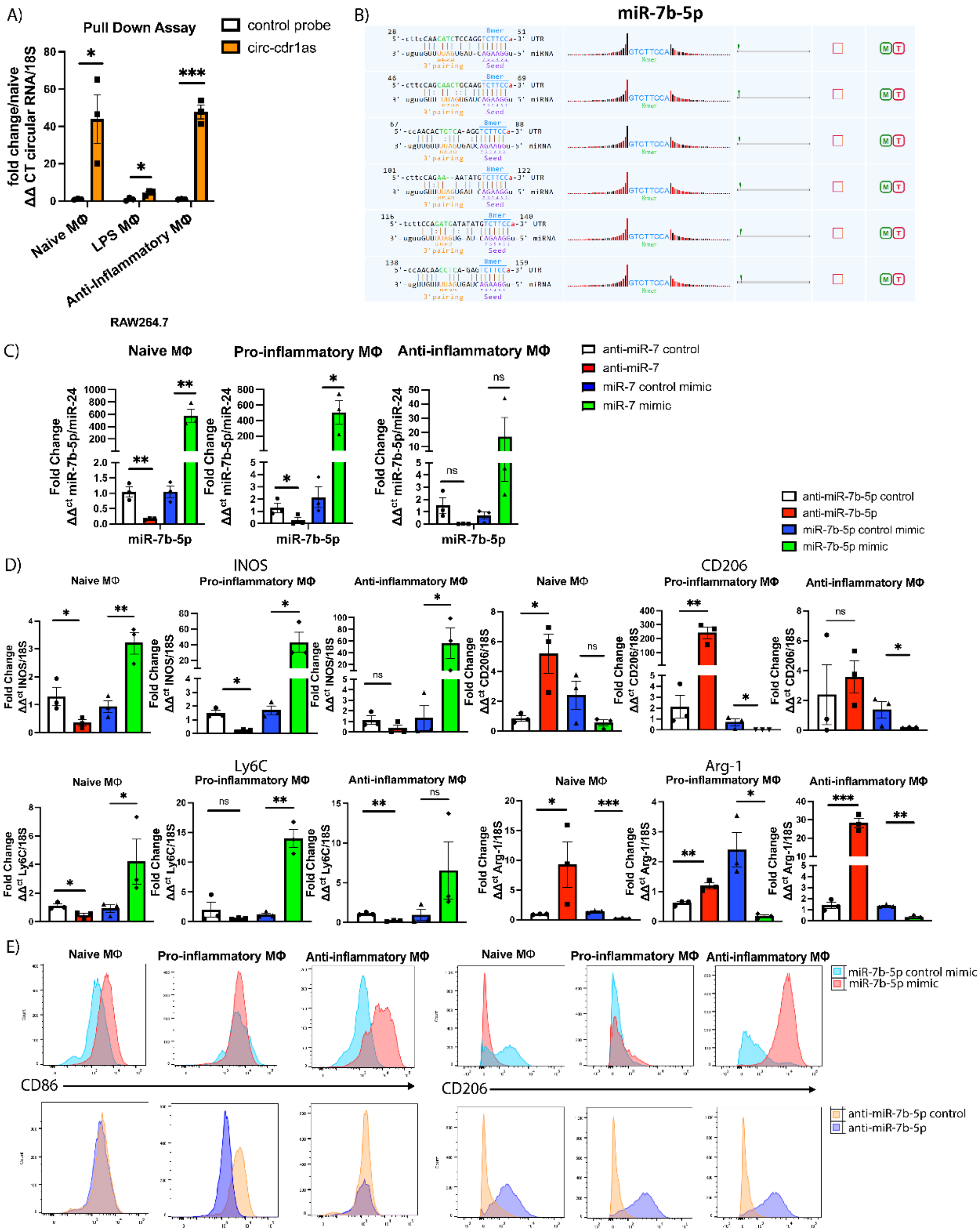


**Supplemental Figure 4 Modification of miR-7b-5p expression alters expression of pro-/anti-inflammatory markers in BMDMs.** (A) Level of circ-cdr1as after pull-down by RT-qPCR in RAW 264.7 cells unpolarized or polarized with LPS or anti-inflammatory cytokines. (Two-sided unpaired t-test). N= 3/group. (B) Map indicating circ-cdr1as and miR-7b-5p binding sites. (C) Validation of miR-7b-5p expression or inhibition by miR-7b-5p mimic or anti-miR-7b-5p, respectively naïve, pro-inflammatory, and anti-inflammatory macrophages. N=3/group. Unpaired t-test. (D) qRT-qPCR analysis of transcriptional changes of macrophage markers in naïve, pro-inflammatory, and anti-inflammatory macrophages treated with miR-7b-5p mimic or anti-miR-7b-5p and their respective controls, normalized to 18S. N=3/group. (Unpaired t-test) Data are mean ± SEM. NS, non-significant, *p<0.05, ** p<0.01, *** p<0.001. (E) Histograms illustrating changes in fluorescence in naïve, pro-inflammatory, and anti-inflammatory macrophages treated with miR-7b-5p mimic or anti-miR-7b-5p and their respective controls. MΦ, macrophages; pro-inflammatory markers: Ly6C, lymphocyte antigen 6 family member C1; INOS, inducible nitric oxide synthase; anti-inflammatory markers: CD206; Arg-1, arginase 1.

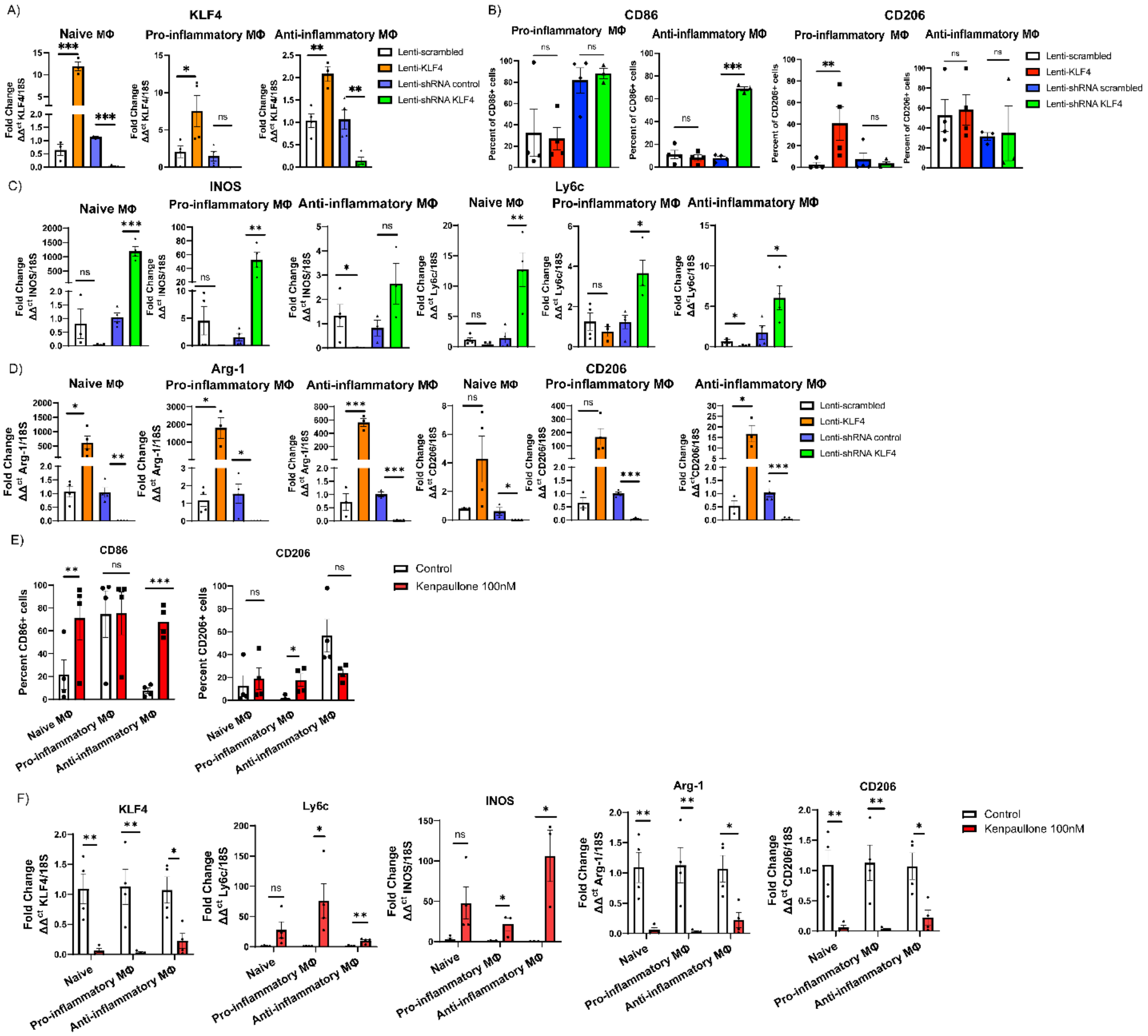

**Supplemental Figure 5 Changes in macrophage marker expression in BMDMs overexpressing KLF4 or knockdown of KFL4.** (A) Validation of KFL4 overexpression expression or inhibition by lentivirus KLF4 or shRNA KLF4 in naïve, pro-inflammatory, and anti-inflammatory macrophages. N=3-4/group (Unpaired t-test). (B) FACS analysis of F4/80/CD86 + cells or F4/80/CD206+ cells in pro-inflammatory and anti-inflammatory macrophages treated with lentivirus KFL4 or shRNA KLF4 and their respective controls. N=3-4/group (One-way ANOVA). RT-qPCR analysis of transcriptional changes of pro-inflammatory macrophage markers (C) or anti-inflammatory macrophage markers (D) in naïve, pro-inflammatory, and anti-inflammatory macrophages treated with lentivirus KFL4 or shRNA KFL4 and their respective controls, normalized to 18S. N=3-4/group (Unpaired t-test). (E) FACS analysis of F4/80/CD86+ or F4/80/CD206+ cells in naïve, pro-inflammatory, and anti-inflammatory macrophages treated with control or kenpaullone 100nM (CDK1/cyclin B, GSK-3β, KLF4 inhibitor) N= 4/group (Unpaired t-test). (F) Validation of KFL4 inhibition by kenpaullone 100nM and RT-qPCR analysis of transcriptional changes of pro/anti- inflammatory macrophage markers (Unpaired t-test) Data are mean ± SEM. NS, non-significant, *p<0.05, ** p<0.01, *** p<0.001. MΦ, macrophages; pro-inflammatory markers: Ly6C, lymphocyte antigen 6 family member C1; INOS, inducible nitric oxide synthase; anti-inflammatory markers: CD206; Arg-1, arginase 1.

### Major Resources Table

#### Animals (in vivo studies)

| Species | Vendor or Source | Background strain | Sex | URL |
|---|---|---|---|---|
| Mouse | Jackson Labs | C57BL/6 #004353 | Male | https://www.jax.org/ |
| Mouse | Jackson Labs | C57BL/6-Tg (UBC-GFP)30Scha/J with a C57BL/6 background | Male | https://www.jax.org/strain/004353 |

#### FACS Antibodies

| Target antigen | Fluoro-phore | Clone | Vendor or Source | Catalog # | Working Concentration | URL |
|---|---|---|---|---|---|---|
| anti-mouse CD16/32 | | S17011E | Biolegend | 156603 | 0.25 µg/100µL | https://www.biolegend.com/fr-ch/products/trustain-fcx-plus-anti-mouse-cd16-32-antibody-17085?GroupID=GROUP20 |
| F4/80 | FITC | BM8 | Thermofisher | 11-4801-82 | 0.5 µg/100µL | https://www.thermofisher.com/antibody/product/F4-80-Antibody-clone-BM8-Monoclonal/11-4801-82 |
| GFP | FITC | FM264G | Biolegend | 338008 | 1 µg/100µL | https://www.biolegend.com/en-us/products/alexa-fluor-488-anti-gfp-antibody-9584 |
| CD206 | APC | C068C2 | Biolegend | 141708 | 0.5 µg/100µL | https://www.biolegend.com/en-us/products/apc-anti-mouse-cd206-mmr-antibody-7425 |
| Ly6C | BV421 | HK1.4 | Biolegend | 128032 | 0.25 µg/100µL | https://www.biolegend.com/en-us/products/brilliant-violet-421-anti-mouse-ly-6c-antibody-8586 |
| CD86 | SB600 | GL-1 | Thermofisher | 63-0862-80 | 0.25 ug/100µL | https://www.thermofisher.com/antibody/product/CD86-B7-2-Antibody-clone-GL1-Monoclonal/63-0862-82 |

### Antibodies for Immunohistochemistry

| Antibody | Host Species or Conjugate | Vendor or Source | Catalog # | Working Concentration | URL |
|---|---|---|---|---|---|
| CD45 | Goat | AF114 | AF114 | 1:200 | https://www.rndsystems.com/products/mouse-cd45-antibody_af114 |
| CD163 | Rabbit | Abcam | AB182422 | 1:30 | https://www.abcam.com/en-us/products/primary-antibodies/cd163-antibody-epr19518-ab182422 |
| CD206 | Goat | R&D systems | AF2535 | 1:50 | https://www.rndsystems.com/products/mouse-mmr-cd206-antibody_af2535?keywords=AF2535 |
| CD68 | Rabbit | Rockland | 600-401-R10 | 1:50 | https://www.rockland.com/categories/primary-antibodies/cd68-antibody-600-401-R10/ |
| GFP | Chicken | Invitrogen | A10262 | 1:100 | https://www.thermofisher.com/antibody/product/GFP-Antibody-Polyclonal/A10262 |
| CD31 | Goat | R&D systems | AF3628 | 1:30 | https://www.rndsystems.com/products/human-mouse-rat-cd31-pecam-1-antibody_af3628?keywords=AF3628 |
| Anti-Actin, α-Smooth Muscle - Cy3™ antibody | Mouse | Sigma | C6198 | 1:1000 | https://www.sigmaaldrich.com/US/en/product/sigma/c6198 |
| Troponin T | Mouse | Invitrogen | MA5-12960 | 1:200 | https://www.thermofisher.com/antibody/product/Cardiac-Troponin-T-Antibody-clone-13-11-Monoclonal/MA5-12960 |
| Wheat Germ Agglutinin, Alexa Fluor™ 488 Conjugate | Alexa Fluor 488 | Thermofisher | W11261 | 1:250 | https://www.thermofisher.com/order/catalog/product/W11261 |
| Donkey anti-rabbit | Alexa Fluor 488 | ThermoFisher | A21206 | 1:100 | https://www.thermofisher.com/antibody/product/Donkey-anti-Rabbit-IgG-H-L-Highly-Cross-Adsorbed-Secondary-Antibody-Polyclonal/A-21206 |
| Donkey anti-mouse | Alexa Fluor 488/555 | ThermoFisher | A21202/ A31570 | 1:100 | https://www.thermofisher.com/antibody/product/Donkey-anti-Mouse-IgG-H-L-Highly-Cross-Adsorbed-Secondary-Antibody-Polyclonal/A-21202 \| https://www.thermofisher.com/antibody/product/Donkey-anti-Mouse-IgG-H-L-Highly-Cross-Adsorbed-Secondary-Antibody-Polyclonal/A-31570 |

| | | | | | |
|---|---|---|---|---|---|
| Donkey anti-goat | Alexa Fluor 488/555 | ThermoFisher | A11055/ A21432 | 1:1000 | https://www.thermofisher.com/antibody/product/Donkey-anti-Goat-IgG-H-L-Cross-Adsorbed-Secondary-Antibody-Polyclonal/A-11055 \| https://www.thermofisher.com/antibody/product/Donkey-anti-Goat-IgG-H-L-Cross-Adsorbed-Secondary-Antibody-Polyclonal/A-21432 |
| Goat anti-chicken | Alexa Fluor 488 | ThermoFisher | A11039 | 1:100 | https://www.thermofisher.com/antibody/product/Goat-anti-Chicken-IgY-H-L-Secondary-Antibody-Polyclonal/A-11039 |

**Enzymes**

| Enzymes | Vendor or Source | Catalog # | Working Concentration | URL |
|---|---|---|---|---|
| collagenase type 2 | Worthington | LS004176 | 250 U/mL | https://www.worthington-biochem.com/products/collagenase?v=705#product-lines |
| collagenase type XI | Worthington | LS004188 | 125 U/mL | https://www.worthington-biochem.com/products/collagenase?v=705#product-lines |
| deoxyribonuclease I | Worthington | LS002060 | 60 U/mL | https://www.worthington-biochem.com/products/deoxyribonuclease-i?v=618 |
| hyaluronidase | Worthington | LS005475 | 60 U/mL | https://www.worthington-biochem.com/products/hyaluronidase#product-lines |

**List of Cytokines for Macrophage Polarization**

| Cytokine | Source or Vendor | Working Concentration | Catalog # | URL |
|---|---|---|---|---|
| IFNγ | R&D systems | 100 ng/mL | 485-MI-100 | https://www.rndsystems.com/products/recombinant-mouse-ifn-gamma-protein_485-mi |
| TNFα | PromoKine | 100 ng/mL | D-63720 | |
| IL-4 | R&D systems | 10 ng/mL | 404-ML-010 | https://www.rndsystems.com/products/recombinant-mouse-il-4-protein_404-ml |
| IL-10 | R&D systems | 10 ng/mL | 217-IL-005 | https://www.rndsystems.com/products/recombinant-human-il-10-protein_217-il |
| TGF-β | R&D systems | 20 ng/mL | 7666-MB-005 | https://www.rndsystems.com/products/recombinant-mouse-tgf-beta-1-protein_7666-mb?keywords=7666-MB-005 |

**List of Primers**

| Species | Gene | Forward Sequence (5'-3') | Reverse Sequence (5'-3') | Manufacturer |
|---|---|---|---|---|
| Mouse | mmu-miR-7b-5p | | | Thermofisher Cat # A25576 Assay ID: mmu483237_mir |
| mouse | mmu-miR-24-3p | | | Assay ID mmu481011_mir Thermofisher Cat # A25576 |
| mouse | Klf4 | GTGGCCCCGGAAAAGAACAG | TTTGCGGTAGTGCCTGGTCA | IDT |
| mouse | Circ-cdr1as | CCTTCACCTCCAAGTCTTCC | TGGAAGATCACGATTGTCTGGA | IDT |
| mouse | 18S | ACG AGA CTC TGG CAT GCT AAC TAG | CGC CAC TTG TCC CTC TAA GAA | IDT |
| mouse | Arg-1 | AAG CCA GGG ACT GAC TAC CTT AAA | TGA TGC CCC AGA TGG TTT TC | IDT |
| mouse | Ly6c | TGC TAT GGA GTG CCA ATT GAG A | GAG CAA TGC AGA ATC CAT CAG A | IDT |
| mouse | CD206 | TTCGGTGGACTGTGGACGAGC A | ATAAGCCACCTGCCACTCCGG T | IDT |
| mouse | INOS | GGGCAGCCTGTGAGACCTT | GCATTGGAAGTGAAGCGTTTC | IDT |

**List of Viruses/miRNA reagents**

| Gene | Product | Catalog # | Vendor or Source |
|---|---|---|---|
| miR-7b-5p | mirVana mRNA mimic cgr mir-7b Assay ID MC26348 | 4464066 | Thermofisher |
| anti-miR-7b-5p | mmu-miR-7b-5p miR-inhibitor Assay ID: MH26348 | 4464084 | Thermofisher |
| mirVana™ miRNA Inhibitor, Negative Control | mirVana™ miRNA Inhibitor, Negative Control | 4464076 | Thermofisher |
| miRNA Mimic, Negative Control | mirVana™ miRNA Mimic, Negative Control | 4464058 | Thermofisher |
| Circ-cdr1as shRNA | Mouse Vector TR30023 pGFP-C-shLenti containing a TCAAGAG loop | TR30023 | OriGene Technologies, Inc., MD, |

| | | | |
|---|---|---|---|
| shRNA scrambled | Mouse Vector TR30023 pGFP-C-shLenti scrambled | TR30023 | OriGene Technologies, Inc., MD |
| Klf4 | NM_010637 Mouse Tagged ORF Clone Lentiviral Par | MR226854L4V | OriGene Technologies, Inc., MD |
| Klf4 control | Mouse Lenti ORF control particles of pLenti-C-mGFP-P2A-Puro | PS100093V | OriGene Technologies, Inc., MD |
| Klf4 shRNA | Klf4 Mouse shRNA Lentiviral Particle | TL313431V | OriGene Technologies, Inc., MD |
| Klf4 shRNA control | Mouse shRNA scrambled Lentiviral Particle | TR30021V | OriGene Technologies, Inc., MD |

**Other (Software)**

| Description | Source | URL |
|---|---|---|
| ImageJ | NIH | https://imagej.net/ij/ |
| Prism | GraphPad | https://www.graphpad.com/ |
| Vevo Lab 5.8.2 | FujiFilm, VisualSonic | https://www.visualsonics.com/application/preclinical/cardiology |
| Illustrator | Adobe | https://www.adobe.com/products/illustrator |
| Excel | Microsoft | https://www.microsoft.com/en-us/microsoft-365 |

**Other (Critical commercial components/Tools)**

| Description | Source | Catalog # | URL |
|---|---|---|---|
| Trichrome Stain (Masson) Kit | Millipore Sigma | HT15 | https://www.sigmaaldrich.com/US/en/product/sigma/ht15 |
| Kenpaullone | MCE | HY-12302 | https://www.medchemexpress.com/Kenpaullone.html |
| miRNeasy Mini Kit | Qiagen | 217004 | https://www.qiagen.com/us/products/discovery-and-translational-research/dna-rna-purification/rna-purification/mirna/mirneasy-kits |
| TaqMan™ Advanced miRNA cDNA Synthesis Kit | Thermofisher | A28007 | https://www.thermofisher.com/order/catalog/product/A28007?SID=srch-srp-A28007 |

| | | | |
|---|---|---|---|
| TaqMan™ Fast Advanced Master Mix | Thermofisher | 4444557 | https://www.thermofisher.com/order/catalog/product/4444557?SID=srch-hj-4444557 |
| Applied Biosystems™ High-Capacity cDNA Reverse Transcription Kit | Thermo Fisher Scientific | 4368814 | https://www.fishersci.com/shop/products/applied-biosystems-high-capacity-cdna-reverse-transcription-kit-4 |
| RNase R | Epicentre Technologies | RNR07250 | https://shop.biosearchtech.com/molecular-biology-enzymes/nucleases/rna-nucleases/ribonuclease-r-%28rnase-r%29/p/NXGENZ-010 |
| Fast SYBR™ Green Master Mix | Thermo Fisher Scientific | 4385612 | https://www.thermofisher.com/order/catalog/product/4385612 |
| VECTASHIELD® PLUS Antifade Mounting Medium | Vector Laboratories, Inc. | H-1900 | https://vectorlabs.com/products/vectashield-plus-antifade?srsltid=AfmBOorIYXYNNd9exbxRkFxiyggAaV_kaT4jD8jvd0w5BZTHqOt14mFV |
| Viral Entry™ Transfection Reagent | Applied Biological Materials Inc. | G515 | https://www.abmgood.com/Transduction-Enhancers.html#ViralEntry |
| Lipofectamine™ RNAiMAX Transfection Reagent | Thermofisher | 13778075 | https://www.thermofisher.com/order/catalog/product/13778075?SID=srch-srp-13778075 |
| HBSS | Gibco | 14170-112 | https://www.thermofisher.com/order/catalog/product/14170112?SID=srch-hj-14170-112 |
| ACK Lysis buffer | Gibco | A10492-01 | https://www.thermofisher.com/order/catalog/product/A1049201?SID=srch-hj-A10492-01 |
| DAPI | Millipore Sigma | 62247 | https://www.thermofisher.com/order/catalog/product/62247?SID=srch-srp-62247 |
| In Situ Cell Death Detection Kit, TMR red (TUNEL) | Roche | 12156792910 | https://www.sigmaaldrich.com/US/en/product/roche/12156792910?srsltid=AfmBOorbqpwAWSYZtciI8PE2ErsGiy8ymxlWF25fxtYSJ2YHxpoTz3Sw |
| $NH_4Cl$ | Stemcell Technologies | 07800 | https://www.stemcell.com/ammonium-chloride-solution.html |

| | | | |
|---|---|---|---|
| NanoDrop 2000 Spectrophotometer | Thermo Fisher Scientific | ND2000 | https://www.thermofisher.com/us/en/home/industrial/spectroscopy-elemental-isotope-analysis/molecular-spectroscopy/uv-vis-spectrophotometry/instruments/nanodrop.html |
| EVOS™ M7000 Imaging System | Thermo Fisher Scientific | AMF7000 | https://www.thermofisher.com/order/catalog/product/AMF7000 |
| T100 PCR thermal cycler | Bio-Rad | 1861096 | https://www.bio-rad.com/en-us/product/t100-thermal-cycler?ID=LGTWGIE8Z |
| StepOnePLus Real-Time PCR system Applied Biosystems 7000 | Thermo Fisher Scientific | 4376600 | https://www.thermofisher.com/us/en/home/life-science/pcr/real-time-pcr/real-time-pcr-instruments/step-one-real-time-pcr-systems.html |
